# Mating-induced Male Death and Pheromone Toxin-regulated Androstasis

**DOI:** 10.1101/034181

**Authors:** Cheng Shi, Alexi M. Runnels, Coleen T. Murphy

## Abstract

How mating affects male lifespan is poorly understood. Using single worm lifespan assays, we discovered that males live significantly shorter after mating in both androdioecious (male and hermaphroditic) and gonochoristic (male and female) *Caenorhabditis*. Germline-dependent shrinking, glycogen loss, and ectopic expression of vitellogenins contribute to male post-mating lifespan reduction, which is conserved between the sexes. In addition to mating-induced lifespan decrease, worms are subject to killing by male pheromone-dependent toxicity. *C. elegans* males are the most sensitive, whereas *C. remanei* are immune, suggesting that males in androdioecious and gonochoristic species utilize male pheromone differently as a toxin or a chemical messenger. Our study reveals two mechanisms involved in male lifespan regulation: germline-dependent shrinking and death is the result of an unavoidable cost of reproduction and is evolutionarily conserved, whereas male pheromone-mediated killing provides a novel mechanism to cull the male population and ensure a return to the self-reproduction mode in androdioecious species. Our work highlights the importance of understanding the shared vs. sex- and species-specific mechanisms that regulate lifespan.

## Introduction

The interplay between the sexes influences an individual’s longevity^1-3^. *Caenorhabditis* female lifespan is shortened after mating through receipt of male sperm and seminal fluid^4^, and separately by exposure to male pheromone^5^. However, previous studies reported contradictory results on how mating influences male lifespan^3,6^. Therefore, whether and how male lifespan is affected by prolonged exposure and interactions with females is largely unknown.

The *Caenorhabditis* genus consists of both androdioecious (male and hermaphroditic) and gonochoristic (male and female) species. In androdioecious species such as *C. elegans,* the population is dominated by hermaphrodites, which reproduce by self-fertilization. Males are usually very rare (less than 0.2% for the standard lab strain N2) and are produced due to spontaneous X chromosome nondisjunction^7,8^. Under stressful conditions, more oocytes experience chromosome non-disjunction, thus androdioecious species periodically undergo explosions of male populations. The existence of males in androdioecious species may reduce inbreeding and facilitate adaptation to changing environments^9^. By contrast, in gonochoristic species such as *C. remanei*, 50% of the population is male, and females and males must mate to reproduce. The mating efficiency of *C. elegans* males is very low compared to *C. remanei* males^8^. Gonochoristic species females secrete pheromones that attract males^10^, and have distinct behaviors during mating compared to hermaphrodites^11,12^. How males in androdioecious and gonochoristic species cope with these different mating situations remains poorly understood. Moreover, the utility of killing females by exposure to male pheromone in gonochoristic populations^5^ is unclear.

Here we report that after mating, *Caenorhabditis* males suffer from germline-dependent shrinking and death, just as in the case of mated *C. elegans* hermaphrodites and *C. remanei* females^4^. However, *C. elegans* males and hermaphrodites have differential sensitivity to male pheromone-dependent toxicity, while *C. remanei* seem immune to this toxicity, and instead use sex-specific pheromones to identify mates. Thus, androdioecious and gonochoristic species differentially utilize pheromone for mating vs hermaphroditic maintenance, while both species suffer the cost of mating through germline-dependent shrinking and death.

## Results

### *C. elegans* males live shorter after mating

*C. elegans* hermaphrodites shrink up to 30% and live 40% shorter after mating^4^. We wondered if males also experience such extreme post-mating changes. Traditional lifespan assays are performed using grouped worms; however, grouped males live shorter than solitary males^13^, which could mask the lifespan shortening effect of mating in males. Therefore, we measured the lifespans of solitary males and single males paired with a single hermaphrodite for 6 days from Day 1 to Day 6 of adulthood. (We used *fog-2(q71)* worms in our assay; *fog-2* males are equivalent to wild-type (N2) males, while *fog-2* hermaphrodites are self-spermless^14^, enabling identification of successful mating.) Mated male lifespan was decreased ~35% compared with the unmated solitary males (Fig. 1A, Table S1), similar to the lifespan decrease of mated hermaphrodites^4^. Also like females, males shrank after 6 days’ mating; by Day 7, the mated males were 10% smaller than the unmated solitary males control (Fig. 1B,C, Table S2).

**Figure 1.**
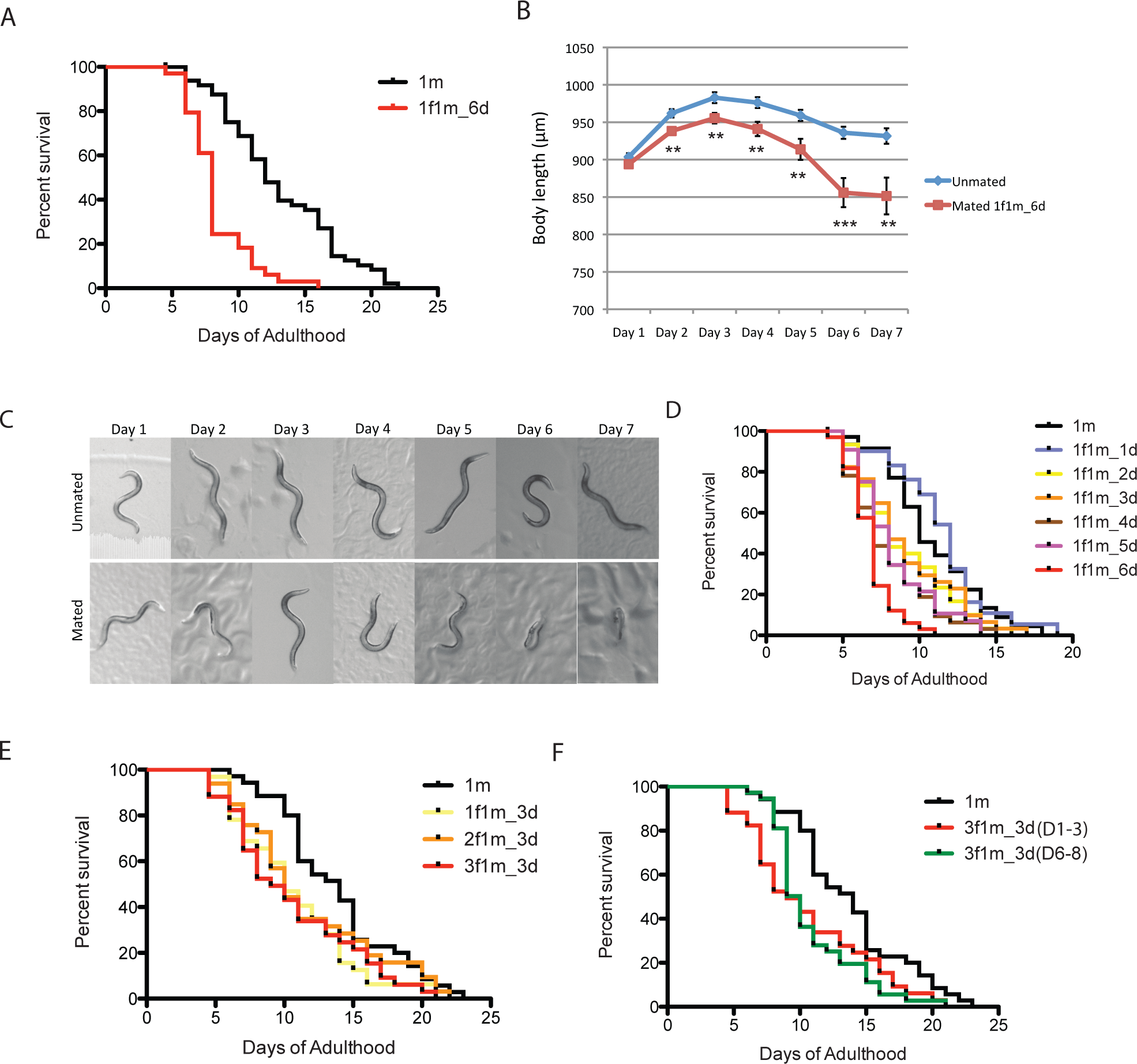
*C. elegans* males shrink and die early after mating. (A) Lifespans of unmated solitary and mated *fog-2(q71)* males. Solitary males: 13.1 ± 0.6 days, n=50; mated males: 8.3 ± 0.4 days, n=34, p<0.0001. Each male was paired with a *fog-2(q71)* hermaphrodite on a single 35mm plate during Day 1-6 of adulthood. Unless noted, all the hermaphrodites used are *fog-2(q71)*. For all the lifespan assays performed in this study, Kaplan-Meier analysis with log-rank (Mantel-Cox) test was used to determine statistical significance. All the lifespan results are included in Table S1. (B) Length of unmated and mated *fog-2* males: t-test, **p<0.01, ***p<0.001. (C) Representative pictures of the same unmated solitary male and male paired with one hermaphrodite from Day 1-Day 6 of adulthood. (D) Male post-mating lifespan decrease is mating duration-dependent: Unmated solitary males: 10.9 ± 0.6 days, n=35; one male and one hermaphrodite mating on Day 1 of adulthood: 11.4 ± 0.6 days, n=31, p=0.3697; mating from Day 1-2: 9.0 ± 0.6 days, n=30, p=0.0325; mating from Day 1-3: 9.1 ± 0.6 days, n=34, p=0.0452; mating from Day 1-4: 7.9 ± 0.5 days, n=32, p=0.0002; mating from Day 1-5: 8.3 ± 0.4 days, n=34, p=0.0006; mating from Day 1-6: 6.8 ± 0.3 days, n=33, p<0.0001. (E) Lifespans of one male paired with different number of hermaphrodites during Day 1-3 of adulthood: solitary unmated males: 13.8 ± 0.7 days, n=35; one male with one hermaphrodite: 10.8 ± 0.6 days, n=32, p=0.0175; one male with two hermaphrodites: 11.6 ± 0.9 days, n=33, p=0.1435; one male with three hermaphrodites: 10.6 ± 0.8 days, n=34, p=0.0147. (F) Lifespans of one male paired with three hermaphrodites for 3 days but at different time of adulthood. Solitary unmated males: 13.8 ± 0.7 days, n=35; mating during Day 1-3 of adulthood: 10.6 ± 0.8 days, n=34, p=0.0147; mating during Day 6-8 of adulthood: 10.8 ± 0.6 days, n=37, p=0.0022.

Males die faster when paired with a hermaphrodite for a longer period: mating with a hermaphrodite for one day did not affect the lifespan of the male, while 2-3 days’ mating shortened male lifespan by 15%, 4-5 days’ mating reduced their lifespan by 25%, and 6 days’ mating increased the reduction to over 35% (Fig. 1D). By contrast, the number of hermaphrodites paired with the single male during mating had much less effect compared to mating duration (Fig. 1E, Fig. S1A,B). The time at which mating occurs within the reproductive period is also not critical for males’ post-mating lifespan decrease; given the same mating duration, males mated with hermaphrodites for the first three days of adulthood had a similar lifespan decrease as those mated with hermaphrodites during Days 6-8 of adulthood (Fig. 1F).

### *C. elegans* males’ post-mating shrinking and death are germline-dependent

We wondered whether pheromone is required for mating-induced death in males. To distinguish pheromone from a direct mating effect, we tested *daf-22(m130)* mutants, which are deficient in ascaroside pheromone biogenesis^15^. Wild-type males still died early post-mating when paired with a *daf-22* hermaphrodite for 6 days (Fig. 2A). Likewise, *daf-22* mutant males lived shorter after 6 days’ mating (Fig. 2B), indicating that the post-mating lifespan decrease in our single-worm pair lifespan assay is due to mating itself rather than pheromone from either sex.

**Figure 2.**
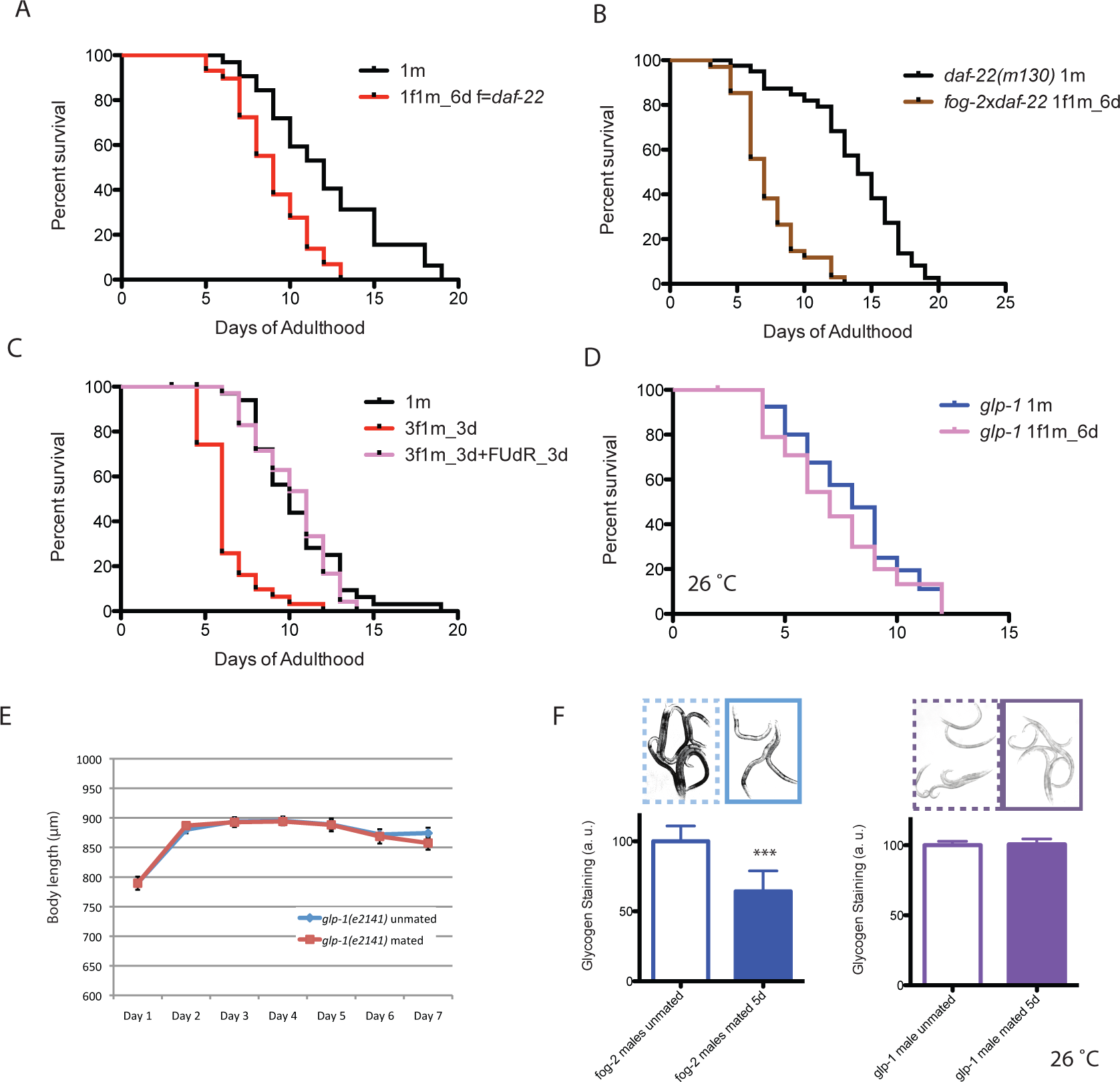
Male post-mating shrinking death is germline-dependent. (A) Lifespans of *fog-2* males mated with *daf-22(m130)* hermaphrodites. Unmated solitary *fog-2* males: 12.1 ± 0.6 days, n=32; mated males: 9.0 ± 0.4 days, n=29, p=0.0001. In the mated group, one *fog-2(q71)* male was paired with one *daf-22(m130)* hermaphrodite from Day 1-Day 6 of adulthood. (B) Lifespans of unmated and mated *daf-22(m130)* males. Unmated solitary *daf-22(m130)* males: 13.8 ± 0.6 days, n=40; mated *daf-22(m130)* males: 7.4 ± 0.4 days, n=34, p<0.0001. In the mated group, one *daf-22(m130)* male was paired with one *fog-2(q71)* hermaphrodite from Day 1- Day 6 of adulthood. (C) FUdR can rescue male post-mating early death. Unmated solitary males: 10.5 ± 0.5 days, n=35; one male with three hermaphrodites for three days: 6.4 ± 0.3 days, n=31, p<0.0001; one male with three hermaphrodites for three days but in the presence of 50 μM FUdR during the three days’ mating: 10.2 ± 0.4 days, n=36, p=0.7086 (compared with unmated solitary group). (D) Lifespans of unmated and mated *glp-1(e2141)* males: unmated solitary *glp-1* males: 8.0 ± 0.4 days, n=40; mated *glp-1* males: 7.2 ± 0.4 days, n=40, p=0.3178. The assay was performed at 26 °C, in mated group, one *glp-1* male was paired with one *fog-2* hermaphrodite from Day 1-6. (E) Length of mated and unmated *glp-1(e2141)* males. (The same population as in Fig. 2D) (F) Glycogen staining of mated and unmated males. Left: mated *fog-2* (wt) males lost over 30% glycogen after 5 days’ mating. *** p<0.001. Right: mated *glp-1* males had a similar amount of glycogen as the unmated *glp-1* males. The staining intensity was normalized to unmated males of each genotype. Representative pictures are shown above the quantitation. Unmated males are framed by dashed lines, and mated males are framed by solid lines.

Elevated germline proliferation is one of the major causes of hermaphrodites’ early death after mating^4^. We wondered whether this killing mechanism is conserved in males. Adult treatment with the DNA replication inhibitor 5-fluorodeoxyruridine (FUdR) has little effect on lifespan and meiosis at low dosage (50 μM)^16^, but rapidly blocks germline proliferation in mated hermaphrodites^4^. When treated with 50 μM FUdR during the three-day mating period, male lifespan was unchanged (Fig. 2C). FUdR treatment also eliminated male post-mating lifespan decrease in our 6 days’ mating regime (Fig. S1C,D). Additionally, lacking the germline prevented both shrinking and death: mating caused neither shrinking nor lifespan decrease in germline-less *glp-1(e2141)* males (Fig. 2D,E, Fig. S1E). These results suggest that germline-mediated post-mating lifespan regulation is conserved between sexes to a large extent.

We have shown previously that osmotic stress resistance correlates well with shrinking in mated hermaphrodites, whereas fat loss does not account for such shrinking^4^. Changes of glycogen levels *in vivo* accurately reflect the osmotic perturbation in the environment^17^; therefore, we measured the glycogen level using iodine staining, and found that mated wild-type worms lost about 30% of the glycogen storage post-mating in a germline-dependent manner (Fig 2F). The mating-induced glycogen storage decrease and subsequent shrinking is conserved between sexes (Fig. S2).

### Vitellogenin dysrégulation contributes to male post-mating death

To further characterize male post-mating death, we performed genome-wide transcriptional analysis of mated and unmated males: we paired a single male with a hermaphrodite for 3.5 days of mating, then picked the males individually from the hermaphrodites on Day 4 for microarray analysis (Fig. S3A). As a control, we collected the same number of age-matched solitary males. 14 genes were significantly up-regulated and 41 were significantly down-regulated (FDR=0%; SAM^18^; Table S3, Fig. 3A). Genes whose expression decreased in mated males include extracellular proteins (*scl-11*, *scl-12, zig-4)* and predicted lipase-related hydrolases *(lips-11, lips-12, lips-13)*. The most enriched gene ontology (GO) categories were ribonucleoside monophosphate biosynthetic/metabolic process and extracellular region for the down-regulated genes, and nutrient reservoir activity and lipid transport for the up-regulated genes (Fig. 3B, Fig. S3B).

**Figure 3.**
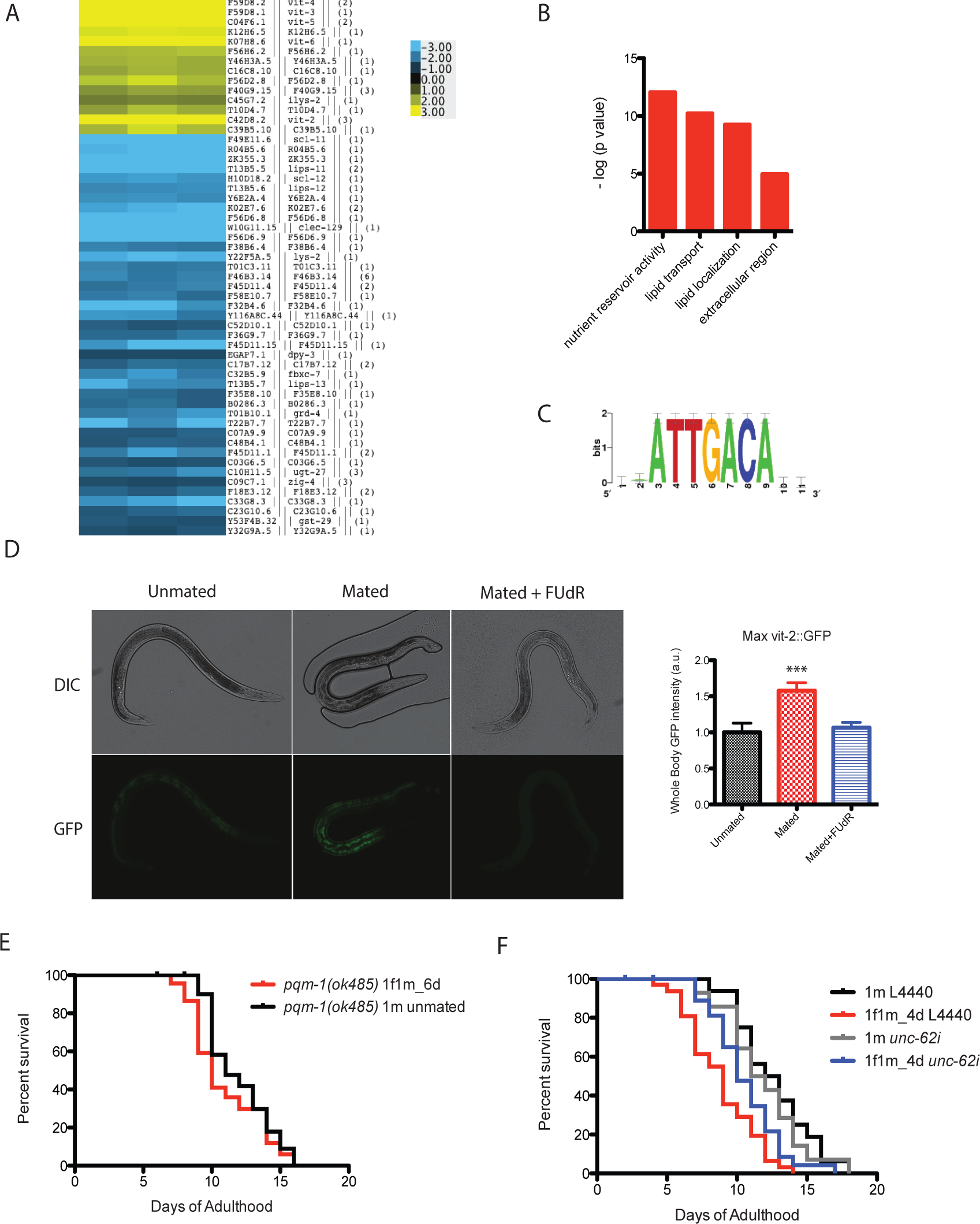
Microarray analysis reveals vitellogenin’s role in male post-mating death. (A) Expression heatmap of genes whose expression is significantly changed in mated males based on SAM analysis. (B) Enriched GO terms for significantly up-regulated genes in mated males. (C) Enriched motif associated with significantly up-regulated genes predicted by RSAT (Regulatory Sequence Analysis Tools). (D) Ectopic expression of VIT-2::GFP in mated males is germline-dependent. 5 days’ mating, pictures were taken on Day 6 of adulthood. Left: images; right: quantification of VIT-2::GFP expression [maximum ± SE (error bars)], a.u., arbitrary units. ***,p<0.001, t-test. (E) *pqm-1(ok485)* mated males have similar lifespans as unmated controls. Unmated solitary *pqm-1(ok485)* males: 11.9 ± 0.5 days, n=25; mated *pqm-1(ok485)* males: 11.0 ± 0.6 days, n=29, p=0.2782. In the mated group, one *pqm-1(ok485)* male was paired with one *fog-2(q71)* hermaphrodite from Day 1- Day 6 of adulthood. (F) *unc-62* RNAi suppresses male post-mating early death. Unmated solitary male on L4440: 12.6 ± 0.7 days, n=25; mated males on L4440: 8.8 ± 0.5 days, n=33, p=0.0001. Unmated males on *unc-62* RNAi: 11.9 ± 0.8 days, n=25; mated males on *unc-62* RNAi: 10.6 ± 0.5 days, n=34, p=0.1249 (compared to unmated males on *unc-62* RNAi).

Surprisingly, vitellogenins (*vit-4, vit-3, vit-5, vit-6, vit-2*), which encode yolk protein precursors made in the female/hermaphrodite intestine for transport into developing oocytes^19^, were the top up-regulated genes in mated males. They were expressed on average 19 times higher in mated males than in solitary unmated males (Table S3). Males normally do not express *vit* genes, as they produce no oocytes. We confirmed our microarray finding using VIT-2::GFP males: mating induced ectopic expression of VIT-2: :GFP, especially in the anterior intestine in males. Such overexpression was germline-dependent (Fig. 3D, S3D). Overproduction of vitellogenins is deleterious for hermaphrodites: vitellogenins accumulate in the head and body of older hermaphrodites^20^; long-lived insulin signaling mutants repress *vit* gene expression^21^; and knockdown of the *vit* genes in wild-type hermaphrodites extends lifespan^21^. The DAE (DAF-16 Associated Element) motif is present in most *vit* genes, which are also Class 2 DAF-16 genes^21^. Thus, we tested the function of PQM-1, the DAE-dependent transcription factor^22^, in male post-mating death. Mated *pqm-1(ok485)* knockout males lived as long as the unmated control (Fig. 3E), suggesting it is important for post-mating death. The binding motif for UNC-62, a master transcription regulator of *vit* genes in hermaphrodites^23^, also emerged in unbiased motif analysis (Fig. 3C). Using RNAi, we found that knocking down *unc-62* was sufficient to rescue the lifespan decrease in mated males (Fig. 3F). Thus, the mis-expression of vitellogenins upon mating contributes to post-mating death in males.

### Mating-induced early death in males is evolutionarily conserved within *Caenorhabditis*

Previously, we showed that *C. remanei* females, like *C. elegans* hermaphrodites, also shrink and die faster after mating^4^, suggesting that the mechanisms are evolutionarily conserved in females. Likewise, we found that male *C. remanei* also lived significantly shorter after mating with a female *C. remanei* for 6 days (Fig. 4A). However, while female death requires successful cross-progeny production, as *C. remanei* males do not induce post-mating death of *C. elegans* hermaphrodites^4^, *C. elegans* males died early when mated with a *C. remanei* female for 6 days (Fig. 4B), suggesting that a component of mating specific and autonomous to the male, rather than a transferred substance or pheromone, is responsible for male death in both species.

**Figure 4.**
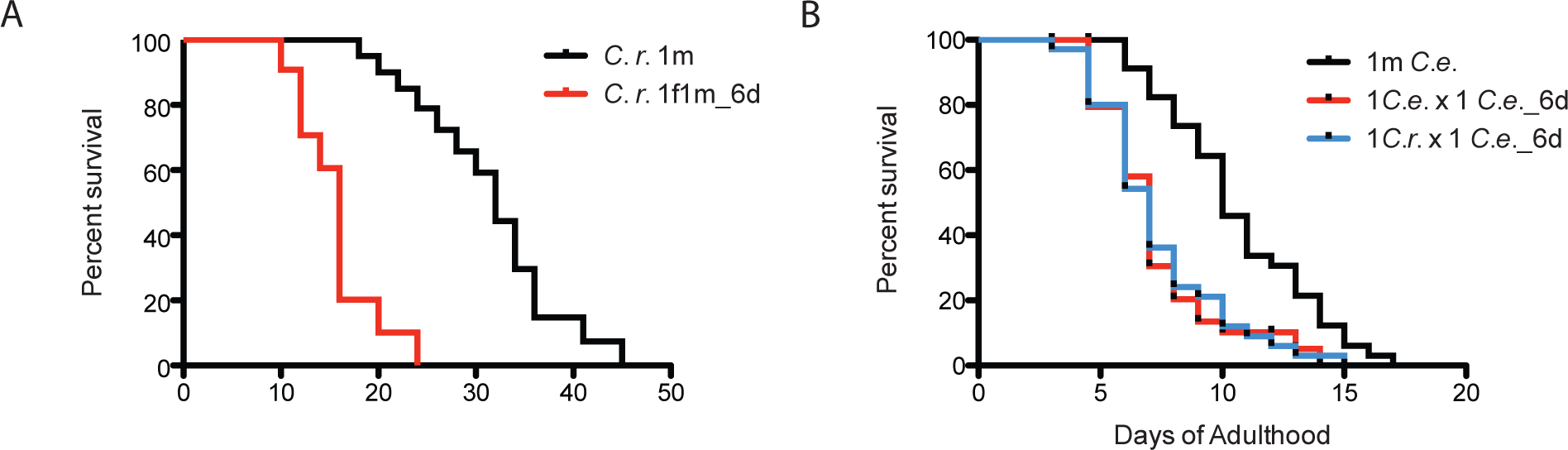
Mating-induced early death in males is conserved. (A) Mated *C. remanei* males also live shorter. Unmated solitary *C. remanei* males: 31.4 ± 1.7 days, n=72; mated *C. remanei* males: 15.7 ±1.2 days, n=28, p<0.0001. In mated group: one *C. remanei* male was paired with one *C. remanei* female from Day 1-Day 6 of adulthood. (B) Lifespans of *C. elegans* males mated with *C. elegans* hermaphrodites and *C. remanei* females. Unmated solitary *C. elegans* males: 10.2 ± 0.6 days, n=35; *C. elegans* males mated with *C. elegans* hermaphrodites: 7.4 ± 0.4 days, n=35, p=0.0001; *C. elegans* males mated with *C. remanei* females: 7.4 ± 0.4 days, n=35, p=0.0003. In mated groups, one *C. elegans* male was paired with either one *C. elegans* hermaphrodite or one *C. remanei* female from Day 16 of adulthood.

### Grouped males also have reduced lifespans in *C. elegans* and *C. remanei*

When male *C. elegans* are housed together, they live shorter compared with solitary males^13^, and the death rate increases with the number of males in a dose-dependent manner^13^ (Fig. 5A). (This might be the reason a previous report failed to report shortened lifespan of males after mating, because grouped males were used as the control^3^.) *C. elegans* male lifespan is very sensitive to male density: just two males together significantly reduced each individual’s lifespan. In a group of eight males, the individual lifespan had a more dramatic 36% decrease compared with the solitary control (Fig. 5A). *C. remanei* male lifespan was also influenced by male density, although to a lesser degree than *C. elegans* males (Fig. 5C). *C. elegans* males tend to form clumps and attempt to mate with each other. By contrast, *C. remanei* males rarely form clumps, having much reduced male-male interaction^13^ (Fig. 5A,C insets). We thought such male-male mating attempts might also lead to post-mating lifespan decrease in a germline-dependent manner as we observed in males mated with females. To test this hypothesis, we placed the grouped males and solitary controls on FUdR plates to inhibit germline proliferation. In the presence of FUdR, grouped *C. remanei* males had no lifespan decrease (Fig. 5D). However, grouped *C. elegans* males still lived significantly shorter (11% decrease compared with solitary control, p=0.0032, Fig. 5B), indicating that a germline-independent factor also contributes to *C. elegans* male lifespan reduction when other males are present.

**Figure 5.**
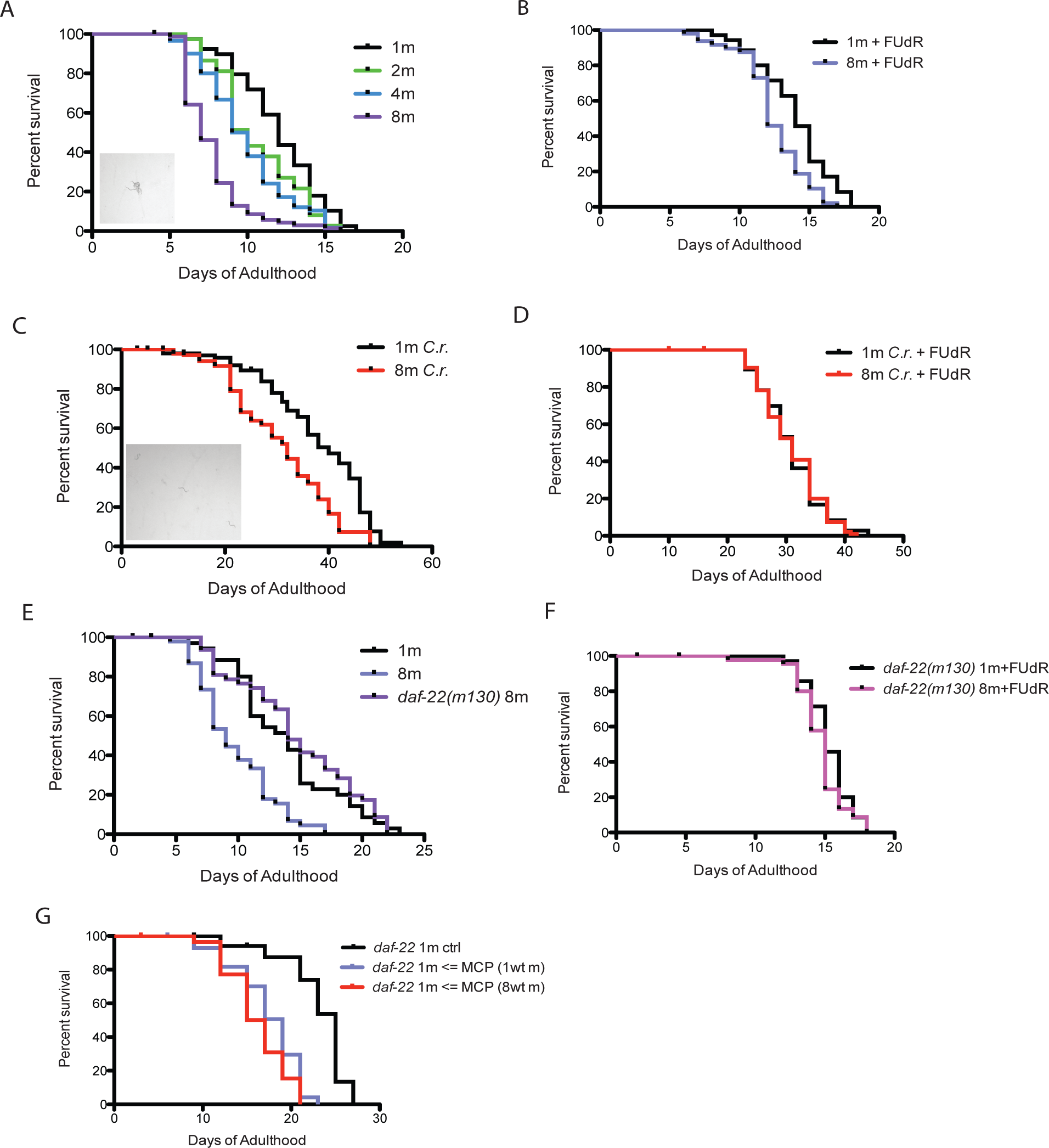
Grouped *C. elegans* males live shorter due to male pheromone. (A) Lifespans of grouped *fog-2(q71)* males. Solitary males: 12.0 ± 0.4 days, n=40; two males: 10.6 ± 0.4 days, n=40, p=0.0397; four males: 9.9 ± 0.4 days, n=60, p=0.0012; eight males: 7.7 ± 0.2 days, n=80, p<0.0001. Inset: clumping of *fog-2* males. (B) Lifespans of grouped *fog-2(q71)* males in the presence of 50μM FUdR. Solitary males: 13.9 ± 0.4 days, n=35; eight males: 12.4 ± 0.3 days, n=48, p=0.0032. (C) Lifespans of grouped *C. remanei* males. Solitary males: 37.9 ± 1.1 days, n=120; eight males: 31.0 ± 0.9 days, n=160, p<0.0001. Inset: *C. remanei* males rarely form clumps. (D) Lifespans of grouped *C. remanei* males in the presence of 50μM FUdR. Solitary males: 30.8 ± 0.9 days, n=45; eight males: 30.8 ± 0.5 days, n=112, p=0.9217. (E) Grouped *daf-22(m130)* males have similar lifespan to solitary wild-type *fog-2* males. Solitary *fog-2* males: 13.8 ± 0.7 days, n=35; eight *fog-2* males: 9.8 ± 0.5 days, n=48, p<0.0001; eight *daf-22(m130)* males: 14.7 ± 0.7 days, n=48, p=0.4039 (compared to solitary males). (F) Lifespans are not different between solitary and grouped *daf-22(m130)* in presence of FUdR. Solitary *daf-22(m130):* 15.3 ± 0.3 days, n=35; eight *daf-22(m130):* 14.7 ± 0.3 days, n=48, p=0.2117. (G) *daf-22(m130)* male lifespans on plates conditioned by wild-type *fog-2* males. MCP: male-conditioned plates. Solitary *daf-22(m130):* 23.0 ± 0.9 days, n=30; *daf-22(m130)* on plates conditioned by one *fog-2* male: 17.3 ± 0.7 days, n=29, p<0.0001; *daf-22(m130)* on plates conditioned by eight *fog-2* male: 16.1 ± 0.6 days, n=30, p<0.0001. Details about male-conditioned plates lifespan assays are included in Methods and Fig. S4B.

### Male pheromone-dependent toxicity leads to reduced lifespan in grouped *C. elegans*

It was shown previously that *C. elegans* hermaphrodites can be killed by male pheromone secreted by grouped males^5^. We wondered whether male pheromone also affects male lifespan. We held 8 *daf-22(m130)* (pheromone-less) males together, and found that they lived as long as the solitary wild-type males, suggesting that male pheromone kills males (Fig. 5E). Grouped *daf-22* males lived just slightly shorter than solitary *daf-22* males (Fig. S4A). The remaining lifespan difference can be explained by germline up-regulation due to mating attempts, since *daf-22* males also formed clumps (Fig. S4A inset), and this lifespan difference was completely eliminated when the experiment was performed in the presence of FUdR (Fig. 5F). Therefore, in grouped *C. elegans* males, early death is due to a combination of germline up-regulation and male pheromone. In fact, males are the victims of their own pheromone: the lifespan of *daf-22* males was significantly reduced when they were maintained on plates conditioned by only one wild-type male (Fig. 5G, S4B), suggesting that *C. elegans* males are extremely sensitive to male pheromone-dependent toxicity.

### *C. elegans* and *C. remanei* have different sensitivity to male pheromone’s toxicity

We wondered whether in a true male/female species, male pheromone-mediated death is also present, and if there are cross-species effects. We confirmed that *C. elegans* hermaphrodites die early when grown on plates conditioned with a large number of *C. elegans* males, as shown previously^5^ (Fig. 6A, 30 males per plate for conditioning). *C. elegans* hermaphrodites also died early when exposed to *C. remanei* male pheromone (Fig. 6A). By contrast, multiple trials of *C*. *remanei* females on male-conditioned plates failed to reveal any sensitivity to either *remanei* or *elegans* male pheromone (Fig. 6B). We then tested the sensitivity of both hermaphrodites and males to low levels of pheromone (8 males per plate for conditioning), and found that *C. elegans* hermaphrodites were not as sensitive to male pheromone as males were (Fig. 6C). By contrast, both *C. remanei* males and females were insensitive to low or high amounts of pheromone (Fig. 6D). Thus, *C. elegans* males are most sensitive to male pheromone-dependent toxicity, *C. elegans* hermaphrodites have intermediate sensitivity, and *C. remanei* appear to be immune to male pheromone toxicity (Fig. S5B).

**Figure 6.**
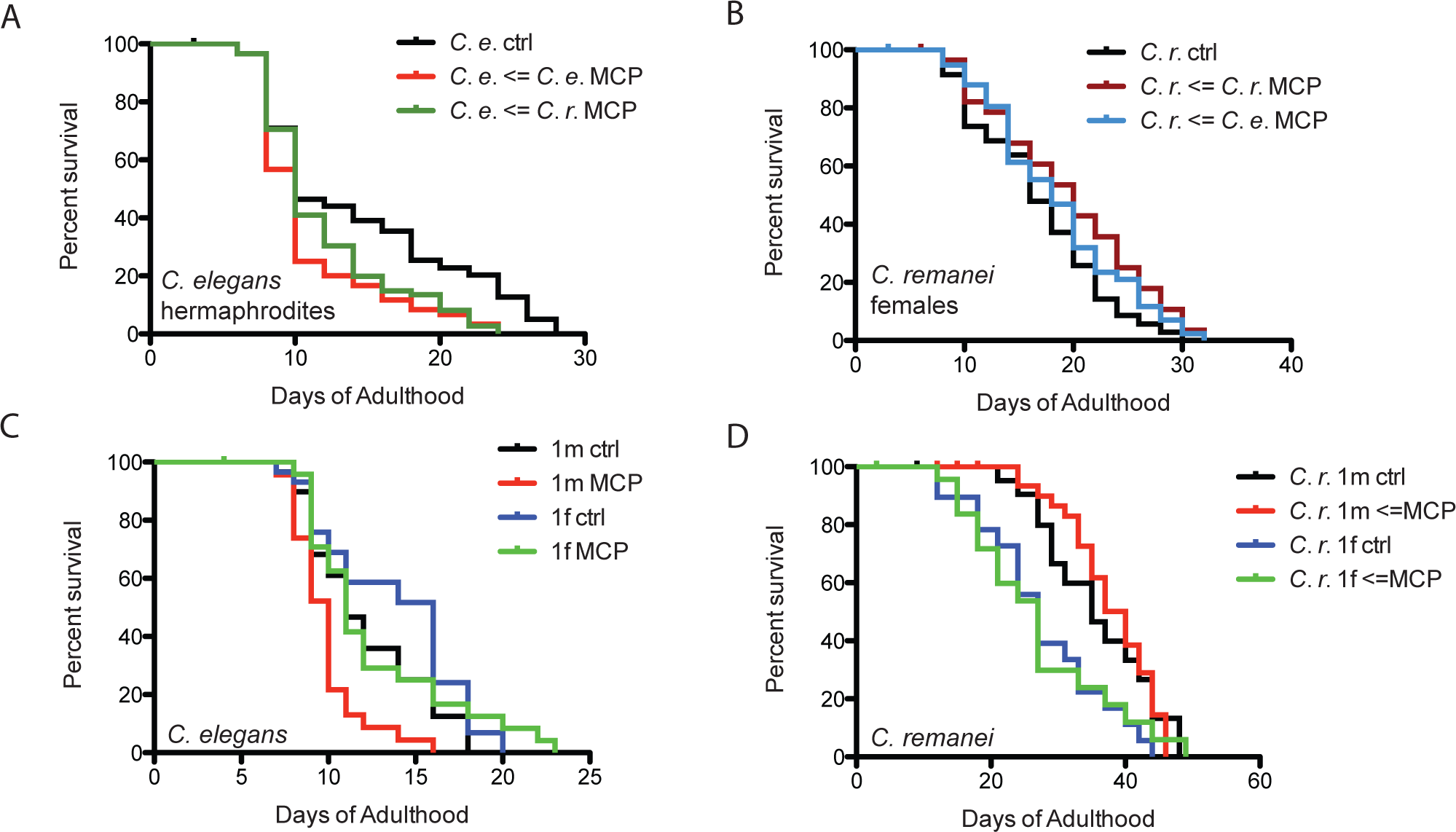
Only *C. elegans* is sensitive to male pheromone’s toxicity. (A) Lifespans of grouped *C. elegans fog-2* hermaphrodites on plates conditioned with 30 males. *fog-2* hermaphrodites control: 14.4 ± 0.8 days, n=90. *fog-2* hermaphrodites on plates conditioned by *fog-2* males: 10.9 ± 0.6 days, n=60, p=0.0004; *fog-2* hermaphrodites on plates conditioned by *C. remanei* males: 11.9 ± 0.5 days, n=90, p=0.0042. (B) Lifespans of grouped *C. remanei* females on plates conditioned by 30 males. *C. remanei* females on control plates: 15.8 ± 0.9 days, n=60; *C. remanei* females on plates conditioned by *C. remanei* males: 19.5 ± 1.3 days, n=30, p=0.0636; *C. remanei* females on plates conditioned by *C. elegans fog-2* males: 18.5 ± 10.9 days, n=60, p=0.1770. (C) Lifespans of solitary *C. elegans fog-2* males and hermaphrodites on plates conditioned by eight *fog-2* males. Solitary *fog-2* males on control plates: 12.1 ± 0.6 days, n=30; solitary *fog-2* males on male-conditioned plates: 9.8 ± 0.4 days, n=28, p=0.0046. Solitary *fog-2* hermaphrodites on control plates: 13.8 ± 0.7 days, n=30; solitary *fog-2* hermaphrodites on male-conditioned plates: 12.6 ± 0.9 days, n=29, p=0.5965. (D) Lifespans of solitary *C. remanei* males and females on plates conditioned by eight *C. remanei* males. Solitary *C. remanei* males on control plates: 35.8 ± 2.0 days, n=34; solitary *C. remanei* males on male-conditioned plates: 37.8 ± 1.2 days, n=34, p=0.8501. Solitary *C. remanei* females on control plates: 27.6 ± 2.2 days, n=24; solitary *C. remanei* females on male-conditioned plates: 27.0 ± 2.5 days, n=30, p=0.8306.

## Discussion

### Germline activation induces deleterious changes that cause males to die

*C. elegans* males and hermaphrodites share many post-mating changes. As we found previously for mated females and hermaphrodites^4^, *Caenorhabditis* males also experience germline-dependent shrinking, glycogen loss, and death after mating. Germline up-regulation also leads to ectopic expression of vitellogenins, which contributes to the post-mating lifespan decrease in males. Previously, these yolk protein precursors were only noted to be expressed in hermaphrodites, since males do not produce oocytes, which normally take up vitellogenins in females. Mating also induces significant overexpression of *vit* genes in hermaphrodites^24^, indicating that vitellogenin expression is closely coupled with mating-induced germline up-regulation in both sexes. Such coupling may be strong enough to overcome the repression of male vitellogenin expression. The striking similarity of germline-dependent post-mating changes in *Caenorhabditis* males and females suggests that this mechanism is largely conserved between sexes, and may represent an unavoidable cost of reproduction as a result of mating.

Germline-dependent lifespan shortening appears to be conserved across species over large evolutionary distances, as it occurs in all *Caenorhabditis* species we tested. Male post-mating death is also conserved beyond the *Caenorhabditis* genus, as *Drosophila* males die earlier after mating, as well (Partridge and Farquhar 1981). To ask whether a similar phenomenon may also present in human males, we examined >2000 years of historical records of ancient imperial China (210 BC-1908 AD), reasoning that emperors should have had the best medical care and highest standard of living available at the time, and extensive notes regarding the emperors’ behavior are available. Although our analysis is limited by the information provided in historical records in ancient China (e.g., other death-contributing factors such as sexually transmitted diseases cannot be ruled out), we censored unnatural deaths (e.g., killed in war) as we would for *C. elegans* studies, and controlled for other factors (e.g., extreme alcohol use). We found that those emperors notorious for lifelong, extremely promiscuous sexual behavior lived 35% shorter than their counterparts (34 ± 2 yrs compared with 52 ± 1 yrs, Fig. 7A, Table S4). Furthermore, analysis of father-son pairs to better control for genetic background and environmental influences (they lived in the same era, therefore had the same standard of living and medical care), still revealed a significant decrease in the lifespan of promiscuous emperors (Fig. 7B-D). While it may seem that any comparison between worms and humans in a germline effect on longevity is highly speculative, it was previously noted that the lifespan of Korean eunuchs was significantly longer than the lifespan of non-castrated men with similar socio-economic status^25^. Together, these results suggest that some aspects of germline-dependent male post-mating death may be evolutionarily conserved.

**Figure 7.**
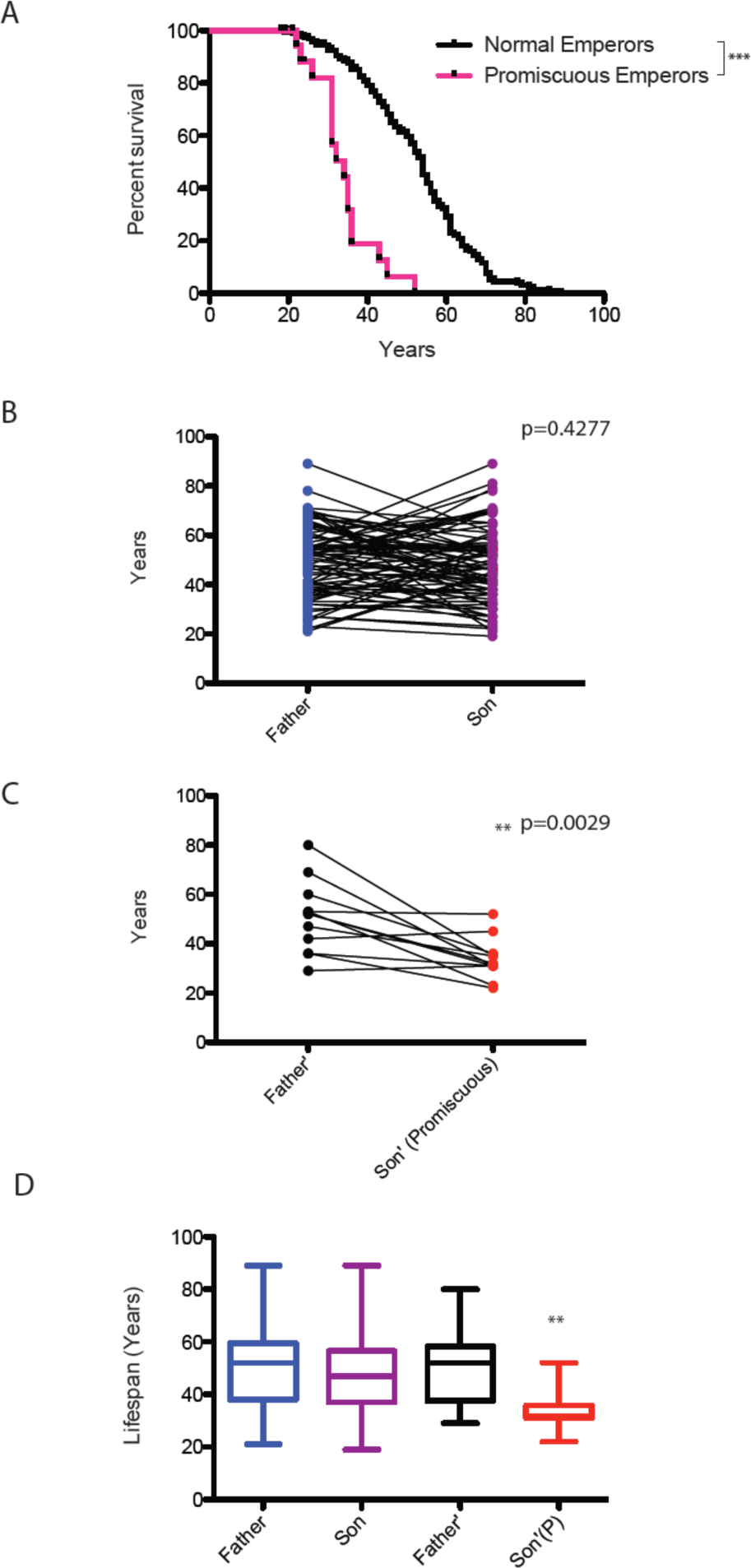
Lifespan analyses of Chinese emperors. (A) Average lifespan of promiscuous Chinese emperors (34 ± 2 yrs, n=21) is 35% shorter than that of nonpromiscuous emperors (52 ± 1 yrs, n=234), p<0.0001. See Methods and Table S4 for detailed rationale, description, and data. (B) There is no lifespan difference between pairs of normal father and son emperors. Father: 49 ± 2 yrs; Son: 48 ± 2 yrs, p=0.4277, n=89, paired t-test. (C) The promiscuous son emperor lives significantly shorter than his father emperor. Father’: 51 ± 4 yrs; promiscuous son: 34 ± 2 yrs; p=0.0029, n=12, paired t-test. The reasons we chose to compare emperor father and son instead of emperor and his brothers are that 1) historical records about emperors’ brothers are much less extensive as those of the emperors themselves; 2) most of these brothers were usually killed by the emperor (or his ally) to ensure his ascendency and to secure his sovereignty. (D) Lifespan summary of (B) and (C).

### Male pheromone-induced killing as a strategy to selectively reduce the male population

In addition to the mating-induced lifespan decrease, *C. elegans* are subject to killing by male pheromone-dependent toxicity, while *C. remanei* are not. Our study shows that androdioecious and gonochoristic species have different sensitivities to male pheromone. The androdioecious species (C. *elegans)* males do not appear to use pheromones as efficiently as chemical messengers to facilitate mating, since they are less able to distinguish hermaphrodites’ pheromone from other species’ female or male pheromone; in fact, *C. elegans* males are even slightly attracted to their own male pheromone, in part explaining their clumping^10^ (Fig. S5A). On the other hand, male pheromone is very toxic to *C. elegans* males. Thus, to *C. elegans* males, pheromones serve primarily as toxins to kill males. By contrast, *C. remanei* (gonochoristic species) males are extremely attracted by pheromone produced by *C. remanei* females, even at a low concentration, and are slightly repelled by male pheromone^10^ (Fig. S5A), but *C. remanei* are immune to both *elegans* and *remanei* male pheromone toxicity (Fig. 6B,D). Thus, the gonochoristic species *C. remanei* uses pheromones primarily as chemical messengers to locate mates. It is also worth noting that such female pheromone-mediated attraction is completely abolished in the presence of male sperm^10^. In *C. elegans*, males are attracted to old, self-spermless hermaphrodites^26,27^, suggesting that pheromone retains the function as a chemical messenger under some circumstances in *C. elegans*. However, due to the presence of self-sperm in the hermaphrodites, *C. elegans* males do not use pheromone as a primary tool to seek young and middle-aged hermaphrodites.

*Caenorhabditis* species might utilize pheromones in such different ways due to their different modes of reproduction. In the androdioecious species *C. elegans*, males are normally rare (0.2%), so the chance that any worm he encounters will be a hermaphrodite is very high; thus, there may be less selection pressure to evolve pheromones as chemical messengers to seek out mates. However, periodically there are explosions of male populations in androdioecious species (e.g., under stressful conditions) to allow outcrossing and ensure genetic diversity^9^. After this period, however, males are more costly to maintain, and there is pressure to return to a primarily hermaphroditic population. It is notable that because *C. elegans* males are XO, rather than XY, males may have no selfish drive to maintain their own chromosomes. From the perspective of species, using male pheromone as a dose-dependent toxin may be an effective way to cull the male population and ensure the species returns to the self-reproduction mode when the stressful condition has passed. Use of the pheromone as a toxin to kill males may have arisen to aid the return to hermaphroditism, which can also be promoted by increased hermaphroditic progeny production and decreased mating rates^28^; these factors could also act in tandem with the selected pheromone-dependent killing of males. Hermaphrodite death at high male pheromone concentration (which would happen extremely rarely in nature) might simply be a rather infrequent result of collateral damage, as the hermaphrodites are less sensitive than males to the toxin. Male-specific culling also occurs in species such as *Drosophila bifasciata*, in which *Wolbachia* infection leads to the killing of male embryos, suggesting that sex ratio can be controlled through male-killing^29^. Mathematical modeling shows that selection in *C. elegans* favors low populations of males^30^, and our model provides a mechanism for how this may be achieved.

By contrast, the preponderance of males in a 50:50 population, as in the case of *C. remanei*, makes the use of pheromone as a toxin less likely, as it would cause too much off-target death to be useful for sperm competition. Our cross-species results suggest that *remanei* male pheromone is toxic to *C. elegans,* but both *C. remanei* males and females are immune to both *elegans* and *remanei* pheromone (Fig. 6A,B). These results also suggest that the toxic effect of pheromone may not be due to the pheromone itself, but rather to a receptor-mediated sensitivity to pheromone that is specific to *C. elegans*, with a greater effect in males than in hermaphrodites. Instead, *C. remanei* pheromone is used to distinguish males from females, an important distinction in 50:50 mixed populations. Like *C. elegans*, the primary mode of sperm competition in *C. remanei* appears to involve seminal fluid transfer of factors that cause the mother to die after producing the father’s progeny, before she has a chance to re-mate^4^, rather than through a pheromone-based mechanism (Fig. 8).

**Figure 8.**
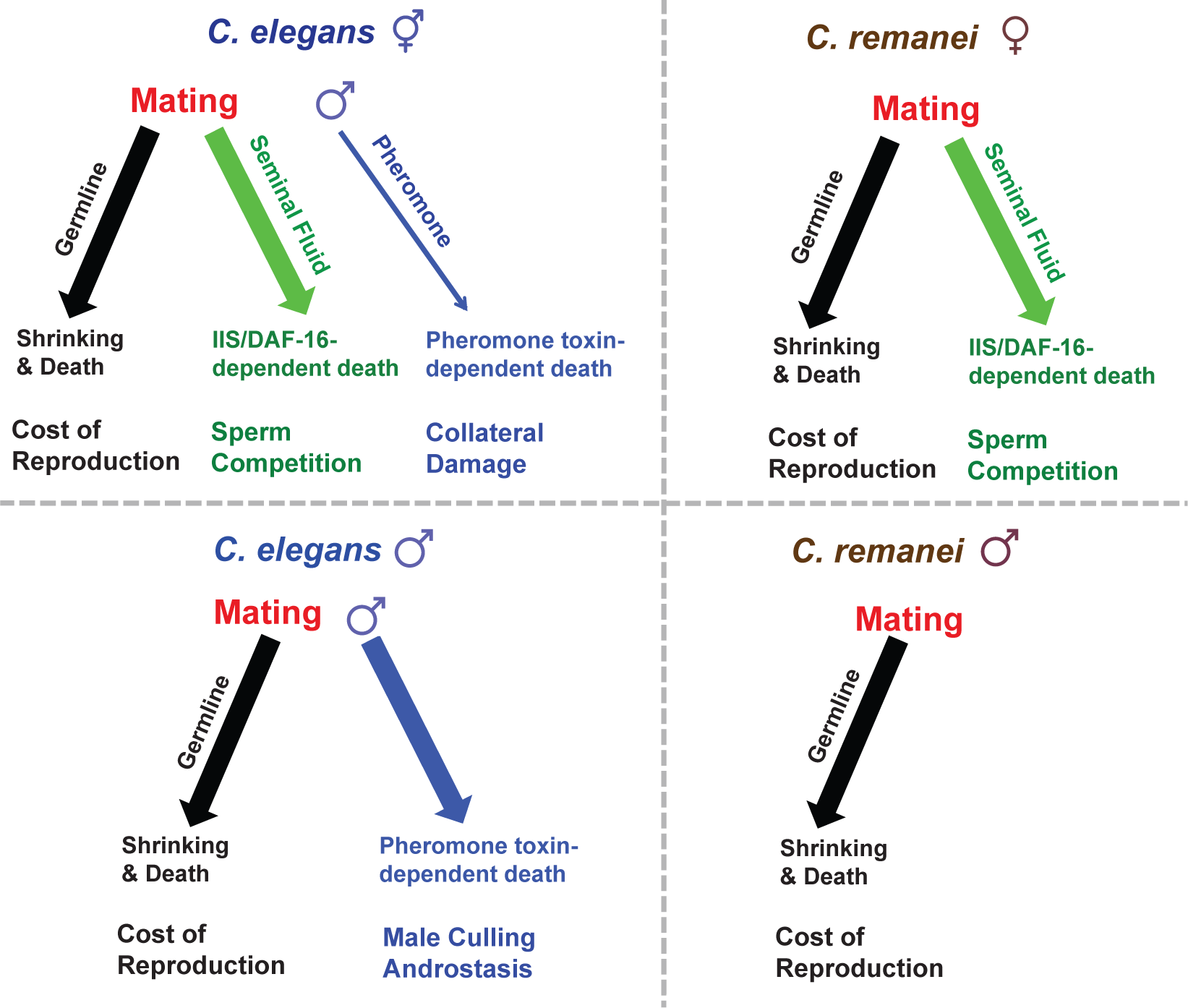
Simplified model of how mating and male pheromone affect lifespan in *C. elegans* hermaphrodites (upper left); *C. remanei* females (upper right); *C. elegans* males (lower left); *C. remanei* males (lower right).

In summary, germline-dependent early death after mating is conserved between sexes and perhaps even across great evolutionarily distances, and is likely due to an unavoidable cost of mating, the result of mated animals ramping up germline proliferation and subsequently exhausting using their own resources as fast as possible to produce the next generation of progeny. The differential use of pheromones as toxins or chemical messengers by males in androdioecious and gonochoristic species demonstrates that they adopt different strategies to compete, mate, and maintain optimal population ratios.

## Acknowledgements

We thank the *Caenorhabditis* Genetics Center (CGC) for strains, Z. Gitai and N. Wingreen for valuable discussions, and members of the Murphy laboratory for critically reading the manuscript. CS is supported by March of Dimes, and AMR by NIH 5T32GM007388-39. CTM is the Director of the Glenn Center for Aging Research at Princeton.

## Author Contributions

C.S., A.M.R., and C.T.M. designed experiments. C.S. and A.M.R. performed experiments. C.S. and C.T.M. wrote the paper.

## Materials and Methods

### Strains

CB4108: *fog-2(q71) V*

CB4037: *glp-1(e2141) III*

DR476: *daf-22(m130) II*

RT130: pwIs23 [vit-2::GFP] (translational fusion)

PB4641: *Caenorhabditis remanei*

### Individual male mating lifespan assays

All the lifespan assays were performed at room temperature (about 20-21°C); except for *glp-1* males’ lifespan assays (performed at 25-26°C). 35mm NGM plates were used for all the experiments in this study. 20 μl of OP50 was dropped onto each plate to make a bacterial lawn of ~10 mm diameter. The next day, one synchronized late L4 male and one late L4 hermaphrodite/female were transferred onto each 35 mm NGM plate. For experiments in Fig. 1E, 1F, S1A-B,D, 2C, multiple L4 hermaphrodites were transferred together with one male. One late L4 male of the same age and genotype was transferred onto the control plates. Except for Fig. 2A, *fog-2(q71)* hermaphrodites were used as the hermaphrodites in the mating assay, because *fog-2* hermaphrodites do not have self sperm, thus allowing us to easily detect successful mating (i.e. eggs and progeny on the plates). We only included males that were able to produce progeny in our analysis. However, for the experiments regarding *glp-1* males, mating on FUdR, and inter-species cross between *C. elegans* males and *C. remanei* females, we included all the males in the analysis. Worms were transferred onto new plates every other day. If the hermaphrodites were lost or bagged, new unmated Day 1 *fog-2* hermaphrodites were added as replacement. Males and hermaphrodites/females were kept together for 6 days (unless noted otherwise in the text); afterwards only males were transferred on to newly seeded plates every 2-3 days. For RNAi experiments in Fig. 3F, synchronized eggs were transferred onto NGM plates with RNAi bacteria, late L4 males were transferred and paired with *fog-2* L4 hermaphrodites onto NGM plates seeded with OP50 (to eliminate the possible effect on mating efficiency for different RNAi treatments). Two days later, males and hermaphrodites were transferred onto fresh plates seeded with corresponding RNAi bacteria and males were maintained on RNAi bacteria thereafter. When lifespan assays were completed, KaplanMeier analysis with log-rank (Mantel-Cox) method was performed to compare the lifespans of different groups.

### Grouped males

35mm NGM plates were used for all the experiments in this study. 20 μl of OP50 was dropped onto each plate to make a bacterial lawn of ~10 mm diameter. The next day, eight synchronized late L4 males were transferred onto each plate. (Two or four males per plate for experiment in Fig. 5A.) One late L4 male of the same age and genotype was transferred onto the control plates. Males were transferred onto fresh plates every two days, when the males were lost or dead, males from other plates were transferred together to make the size of the group stable.

### Male-conditioned plates (MCP) setup

Male-conditioned plates for lifespan assays were prepared as previously described^5^. Briefly, 60 μl of OP50 was dropped onto each 35mm NGM plate to make a bacterial lawn of ~25 mm diameter. Young Day 1 wild-type males (*fog-2* males) were transferred onto each plate. Two days later, they were removed and worms for lifespan assays were immediately transferred onto these male-conditioned plates. These male-conditioned plates were being prepared throughout the course of the lifespan assays (Fig. S4B). For the experiments in Fig. 6A,B, 30 males were used for each conditioning plate. For experiments in Fig. 6C,D, 8 males were used for conditioning and for the experiment in Fig. 5G, only 1 male was used for conditioning for each plate.

### Body size measurement

Images of live males on 35mm plates were taken daily for the first week of adulthood with a Nikon SMZ1500 microscope. Image J was used to analyze the body size of the worms. The middle line of each worm was delineated using the segmented line tool and the total length was documented as the body length of the worm. T-test was performed to compare the body size differences between groups of males in the same day.

### FUdR experiment

FUdR was added to the NGM media to the final concentration of 50 μM. Late L4 males and hermaphrodites were transferred onto NGM+FUdR plates seeded with OP50. Worms were transferred every two days, and were kept on FUdR plates for different period of time (3 days, 6 days or lifetime as indicated by text).

### Glycogen staining

Glycogen staining was performed according to a well-described protocol^17^. Mating of males was set up as previously described. Right before staining, live males of the same group were picked into a M9 droplet with 1M sodium azide on a 3% agarose pad. Immediately after the liquid was dry, the pad was inverted over the opening of a 50g bottle of iodine crystal chips (Sigma) for 1 minute. After the color stained by iodine vapor on the pad disappear (non-specific staining), the worms were immediately imaged by a Nikon microscope. Due to uncontrollable differences, it is hard to compare the staining performed at different times. Thus, worms from the groups of comparison were mounted onto the same pad (separate M9 droplet if there is no visible difference). Image J was used to compare the mean intensity of iodine staining after the background was subtracted. T-test was performed to compare the staining between different groups (on the same pad).

### GFP intensity quantification

10-20 worms of each group were imaged by Nikon Ti. Image J was used to measure the mean and the maximum GFP intensity of the whole body area. T-test analysis was performed to compare the GFP intensity of different groups of worms.

### Mated males microarrays

We paired a single male with a *fog-2* hermaphrodite for about 3.5 days of mating, then picked the males individually from the hermaphrodites on Day 4 for microarray analysis. As a control, solitary males were collected at the same time. About 150 males (on 150 individual 35mm plates) were collected for each condition and replicate. Three biological replicates were performed. RNA was extracted by heat-vortexing method. Two-color Agilent microarrays were used. The detailed steps and analysis were performed according to a previous report^31^.

### Pheromone chemotaxis assay

This assay was modified from a previous assay^10^. 10 Day 1 virgin *C. remanei* or *C. elegans* hermaphrodites were put in 100 μl of M9 buffer at room temperature overnight with shaking. 100 males of either *C. elegans* or *C. remanei* were put in 100 μl of M9. The supernatant solutions were then taken for pheromone chemotaxis assay. 60 mm NGM plates (no food) were used for the chemotaxis assay. Two destination spots (supernatant and M9 control) were separated by about 45 mm, the distance from the origin spot to either destination spot is 30mm. Two 1 μl drops of 1M sodium azide were first applied to the destination spots. When dry, a drop of 1 μl M9 or supernatant was separately added onto the destination spots. Then, over 10 young adult (Day 2) males were placed at the origin spot (try to transfer as little bacteria as possible). After 60 minutes, the paralyzed male worms were scored based on their location. The chemotaxis index was calculated as: (#worms at supernatant destination - #worms at control destination)/(#total worms - #worms at origin).

### Analysis of Lifespans of Emperors in Imperial China

In ancient China, agriculture was the main source of the country’s wealth. The development of agriculture began in the Neolithic Era (10,000 BC), followed by improvements in the Bronze Age (1000 BC). Late in the Warring states eras (771-221 BC), new iron tools were widely adopted, which revolutionized agriculture in China. Ancient China’s economy depended heavily if not solely on agriculture.

Qin Shi Huang (#1 on the list below) was the first emperor to unify China. By that time, agriculture had already been well developed and the basic structure and the quality of civilization did not change much until the late 1800s. Emperors had the best standard of living and medical care at the time, and the living conditions of emperors in Imperial China (220 BC -1911) remained relatively similar (i.e., the best of agricultural civilization) over this period of 2000 years.

To perform our lifespan analysis analogously with the approach we use to assess worm lifespan, we only included emperors who were over 18 years old when they died and those who reigned over 1 year, in order to exclude the cases of puppet emperors (Table S4). Those rows marked by grey on the list indicate that the emperor’s death is unnatural (killed in a war, rebellion, etc); we censored these emperor at the time of death, analogously to how we would censor worms who died unnaturally or disappeared during a lifespan assay. Those highlighted in yellow are emperors with extremely promiscuous sexual behaviors, as documented by official historical records. Those labeled by shaded yellow means they were considered promiscuous but died unnaturally.

The average lifespan of promiscuous emperors was 34 years, which is 35% shorter than the normal emperors’ lifespan (52 years) (Fig. 7, Table S4). It should be noted that these promiscuous emperors were also noted to indulge in excessive alcohol consumption; however, other emperors who were well-known for their lifelong alcohol indulgence were not short-lived (Examples are Yuan Tai Zong #216 on the list, died at 56; Yuan Shi Zu #219 died at 79).

Another case worth noting is Song Gao Zong (#178), who was originally fertile but is reported to have become infertile when he fled south after defeat by his enemies. By the time he reestablished his dynasty in southern China, he was only 24, but was reportedly no longer capable of reproduction; he died at the age of 81. His case may suggest the link between germline signal and lifespan, perhaps in the same manner as the suggested lifespan extension of Korean eunuchs documented by Min, et al. (2012).

### Analysis of father-son comparisons

To better control for genetic background and environmental influences, the lifespans of father and son emperors was compared. The reasons we chose to compare father and son instead of emperor and his brothers are twofold: 1) historical records about emperors’ brothers are much less extensive as those of the emperors themselves; 2) most brothers were killed by the emperor (or his ally) to ensure his ascendency and to secure his sovereignty.

## Main Figure Legends

### Abbreviations and nomenclature in the paper

*C. e.*: *C. elegans*
*C. r.*: *C. remanei*
1f1m_6d: “f” stands for hermaphrodite/female, “m” stands for male, the number before f/m suggests the amount of worms on the same 35mm plate. “6d” means mating for 6 days.
AAA x BBB: hermaphrodites/females of genotype AAA are mated with males of genotype BBB. (male is always listed after the “x”)
MCP: male-conditioned plates

## Supplementary Figures

**Fig. S1.**
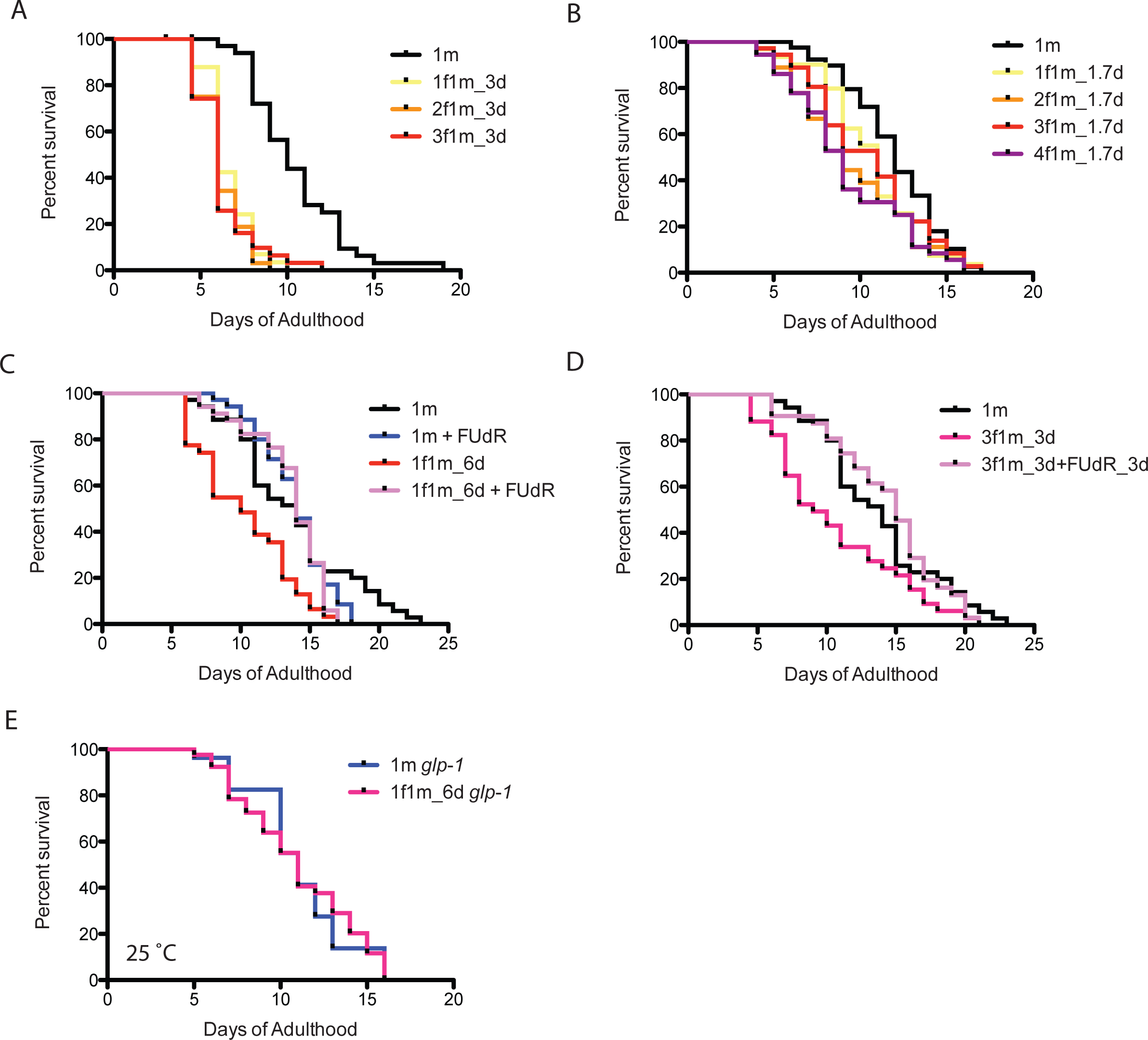
How mating affects male lifespan. (A) Lifespans of one male paired with different number of hermaphrodites during Day 1-3 of adulthood: solitary unmated males: 10.5 ± 0.5 days, n=35; one male with one hermaphrodite: 6.6 ± 0.2 days, n=33, p<0.0001; one male with two hermaphrodites: 6.3 ± 0.2 days, n=32, p<0.0001; one male with three hermaphrodites: 6.4 ± 0.3 days, n=31, p<0.0001. (B) Lifespans of one male paired with different number of hermaphrodites for the first 1.7 days of adulthood: solitary unmated males: 12.0 ± 0.4 days, n=40; one male with one hermaphrodite: 10.6 ± 0.5 days, n=32, p=0.0824; one male with two hermaphrodites: 9.7 ± 0.6 days, n=37, p=0.0435; one male with three hermaphrodites: 10.4 ± 0.6 days, n=36, p=0.1575; one male with four hermaphrodites: 9.3 ± 0.6 days, n=36, p=0.0041. (C) FUdR can rescue male post-mating early death. Unmated solitary males: 13.8 ± 0.7 days, n=35; one male with one hermaphrodite for six days: 10.3 ± 0.6 days, n=31, p=0.0006; solitary male in the presence of 50 μM FUdR: 13.9 ± 0.4 days, n=35, p=0.4079 (compared to unmated solitary group). One male mating with one hermaphrodites for 6 days in the presence of FUdR: 13.6 ± 0.5 days, n=34, p=0.3992 (compared to unmated solitary group). (D) Lifespans of males paired with three hermaphrodites for 3 days with FUdR: unmated solitary males: 13.8 ± 0.7 days, n=35; one male with three hermaphrodites for three days: 10.6 ± 0.8 days, n=34, p=0.0147; one male with three hermaphrodites for three days but in the presence of 50 μM FUdR during the three days’ mating: 14.3 ± 0.7 days, n=32, p=0.8740 compared with unmated solitary group. (E) Lifespans of unmated and mated *glp-1(e2141)* males: unmated solitary *glp-1* males: 11.1 ± 1.0 days, n=27; mated *glp-1* males: 11.1 ± 0.5 days, n=43, p=0.9149. The assay was performed at 25 °C, in mated group, one *glp-1* male was paired with one *fog-2* hermaphrodite from Day 1-6.

**Fig. S2.**
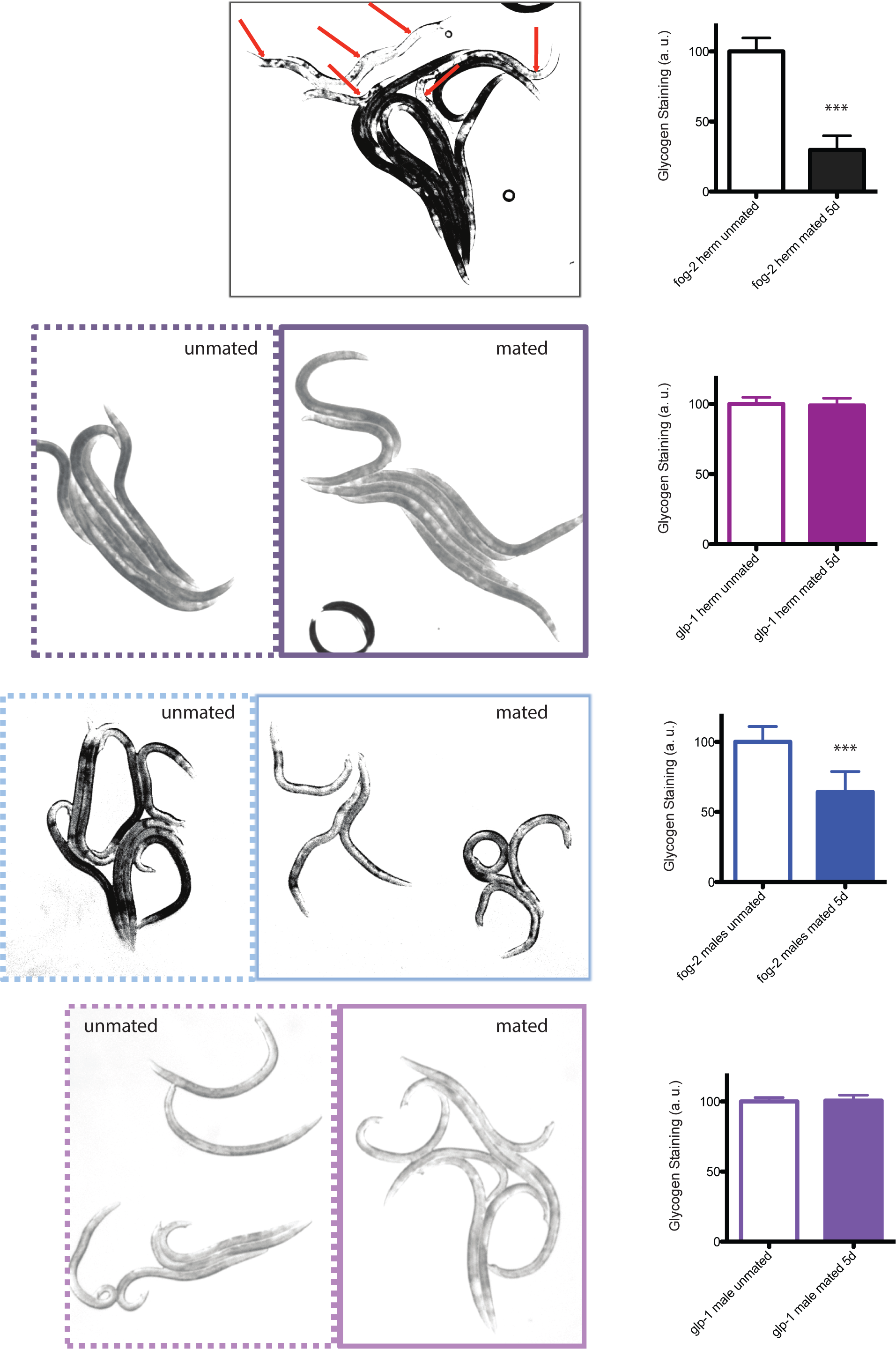
Glycogen staining of mated vs unmated hermaphrodites and males. Left: representative pictures of iodine staining of worms. Unmated worms are framed by dashed lines, whereas mated worms are framed by solid lines. In the first picture, mated and unmated *fog-2* hermaphrodites were mixed together, with red arrows pointing to mated *fog-2* hermaphrodites. Worms were mated from Day 1 – Day 5 and were imaged on Day 5. Right: quantitation of iodine staining. The intensity of mated worms was normalized to unmated control of the same genotype. Mated *fog-2* hermaphrodites have 30% glycogen compared to unmated *fog-2* hermaphrodites of the same age (p<0.0001). Mated *glp-1* hermaphrodites have 99% glycogen compared to unmated *glp-1* hermaphrodites control (p=0.6070). Mated *fog-2* males have 64% glycogen compared to unmated *fog-2* males of the same age (p<0.0001). Mated *glp-1* males have 101% glycogen compared to unmated *glp-1* males control (p=0.7107). Error bars represent SD. ***, p<0.0001, t-test.

**Fig. S3.**
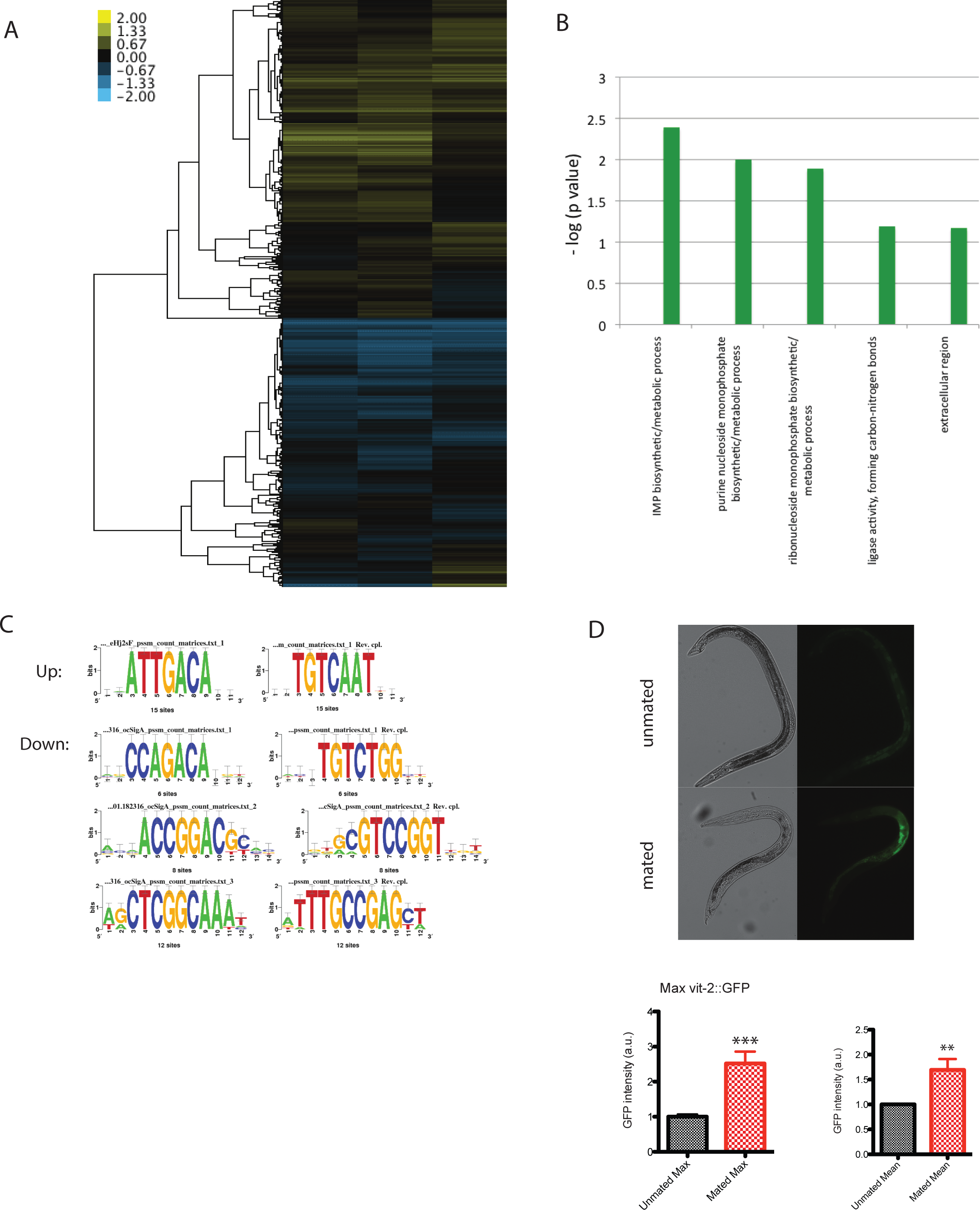
Microarray analysis of mated males. (A) Expression heat map of clustered mated males vs unmated males. Individual males were paired with one hermaphrodite for 3.5 days and collected on Day 4 for microarrays. (B) Enriched GO terms for significantly down-regulated genes in mated males. (C) Enriched motifs in promoter region (1kb upstream of TSS) of significantly up- and down-regulated genes using (RSAT) Regulatory Sequence Analysis Tools (www.rsat.eu). (D) VIT-2::GFP expression in males increases significantly after mating. Upper: DIC and GFP images; Lower: GFP intensity quantitation, left: max ± SE (error bars); right: mean ± SE (error bars), a.u., arbitrary units. **, p<0.01, t-test.

**Fig. S4.**
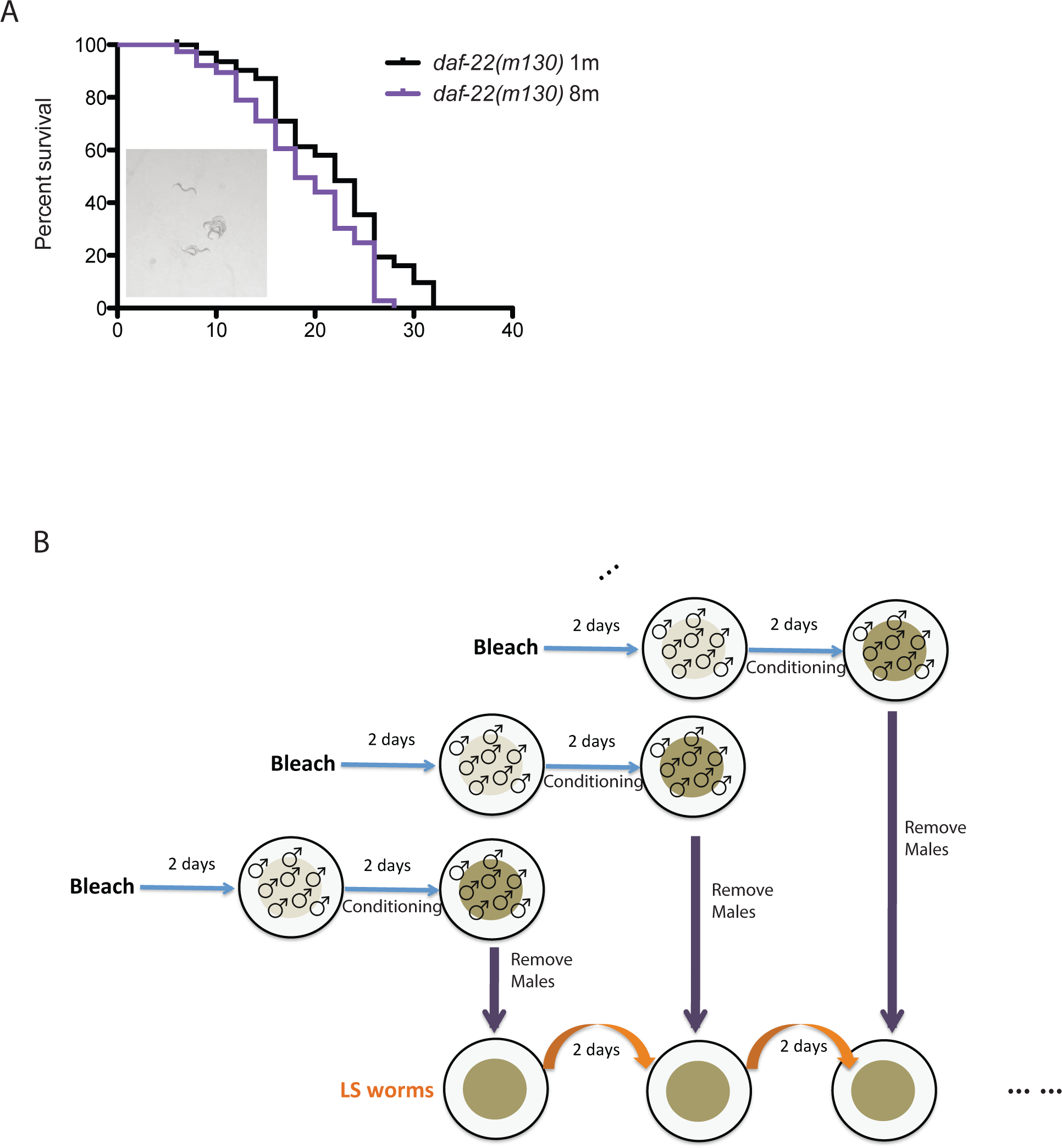
Male pheromone and male-conditioned plates (MCP). (A) Lifespans of grouped *daf-22(m130)* males. Solitary males: 21.7 ± 1.2 days, n=32; eight males: 18.8 ± 1.0 days, n=38, p=0.0394. Inset: *daf-22(m130)* males also form clumps. (B) Schematic illustration of how lifespan assays on male-conditioned plates were performed. Detailed description is included in Methods.

**Fig. S5.**
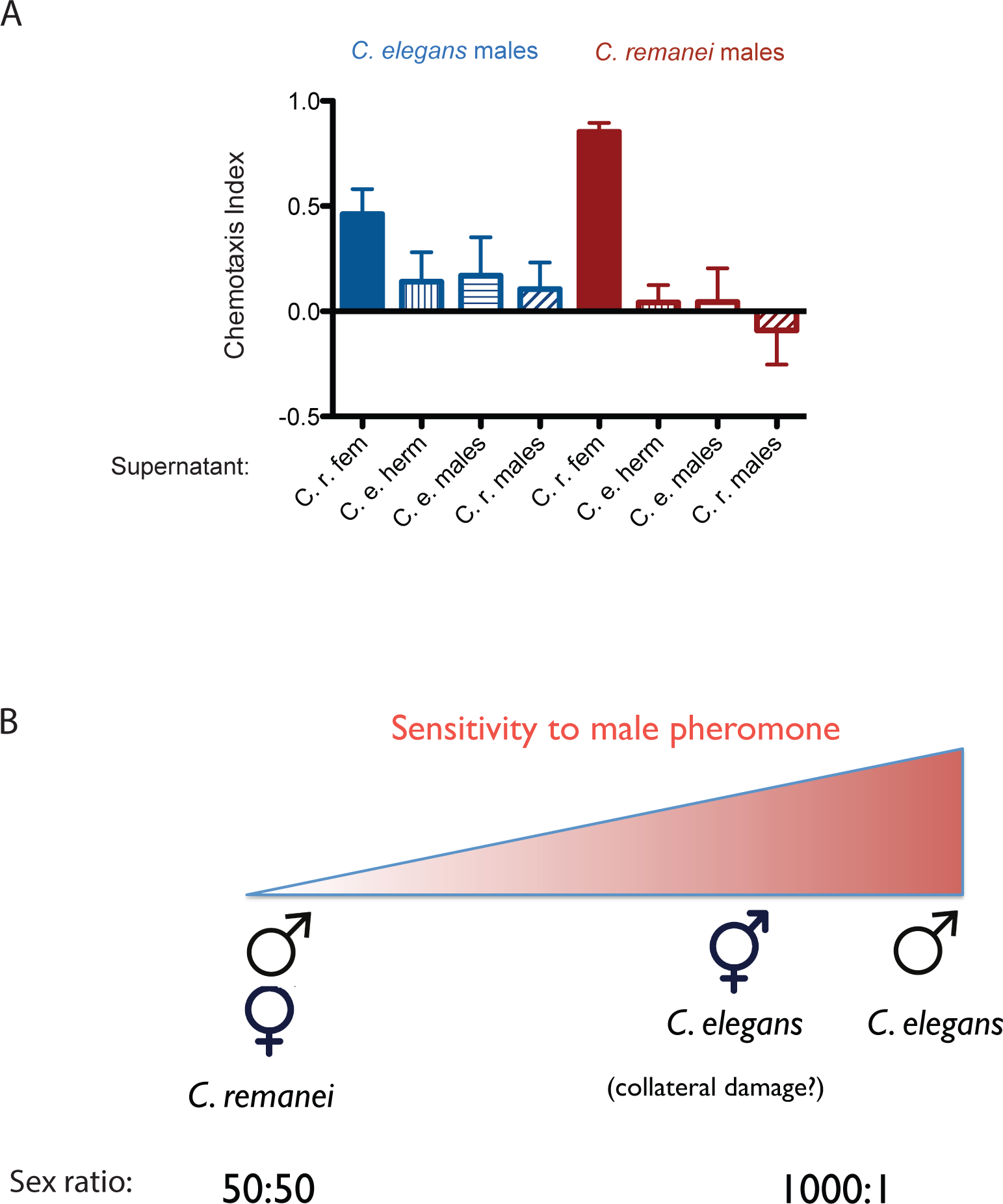
Male chemotaxis to different pheromones. (A) Supernatant solutions from *C. elegans* males, *C. remanei* males, *C. elegans* N2 hermaphrodites, and *C. remanei* females are used to do the chemotaxis assay. See Methods for detailed description. *C. e*. males to supernatant of *C. r*. females: Chemotaxis Index (CI) is 0.46 ±0.11 (mean ± SEM, n=12 [plates]); *C. e*. males to supernatant of *C. e*. hermaphrodites: CI = 0.14 ± 0.13 (n=10); *C. e*. males to supernatant of *C. e*. males: CI = 0.17 ± 0.17 (n=12); *C. e*. males to supernatant of *C. r*. males: CI = 0.11 ± 0.12 (n=11); *C. r*. males to supernatant of *C. r*. females: CI = 0.85 ± 0.04 (n=12); *C. r*. males to supernatant of *C. e*. hermaphrodites: CI = 0.04 ± 0.08 (n=12); *C. r*. males to supernatant of *C. e*. males: CI = 0.04 ± 0.15 (n=12); *C. r*. males to supernatant of *C. r*. males: CI = −0.09 ± 0.16 (n=12). (B) Toxicity scale of sensitivity to male pheromone.

## Supplementary Tables

**Table S1.** Lifespan assays summary.

| Genotype/condition | Mean LS ± std. error | % change | p value | N | Related Figure |
| --- | --- | --- | --- | --- | --- |
| **Experiment 1** |  |  |  |  |  |
| 1m *fog-2* | 13.1 ± 0.6 | -- | -- | 50 | Fig. 1A |
| 1f1m_6d *fog-2* | 8.3 ± 0.4 | -37% | <0.0001 | 34 | Fig. 1A |
| 1f1m_1.5d *fog-2* | 11.9 ± 0.6 | -9% | 0.0988 | 38 | |
| 2m *fog-2* | 11.4 ± 0.5 | -13% | 0.0149 | 48 | |
| 4m *fog-2* | 9.3 ± 0.6 | -29% | <0.0001 | 48 | |
| **Experiment 2** |  |  |  |  |  |
| 1m *fog-2* | 10.9 ± 0.6 | -- | -- | 35 | Fig. 1D |
| 1f1m_1d *fog-2* | 11.4 ± 0.6 | +5% | 0.3697 | 31 | Fig. 1D |
| 1f1m_2d *fog-2* | 9.0 ± 0.6 | -17% | 0.0325 | 30 | Fig. 1D |
| 1f1m_3d *fog-2* | 9.1 ± 0.6 | -17% | 0.0452 | 34 | Fig. 1D |
| 1f1m_4d *fog-2* | 7.9 ± 0.5 | -28% | 0.0002 | 32 | Fig. 1D |
| 1f1m_5d *fog-2* | 8.3 ± 0.4 | -24% | 0.0006 | 34 | Fig. 1D |
| 1f1m_6d *fog-2* | 6.8 ± 0.3 | -38% | <0.0001 | 33 | Fig. 1D |
| **Experiment 3** |  |  |  |  |  |
| 1m *fog-2* | 13.8 ± 0.7 | -- | -- | 35 | Fig. 1E |
| 1f1m_3d *fog-2* | 10.8 ± 0.6 | -22% | 0.0175 | 32 | Fig. 1E |
| 2f1m_3d *fog-2* | 11.6 ± 0.9 | -16% | 0.1435 | 33 | Fig. 1E |
| 3f1m_3d *fog-2* | 10.6 ± 0.8 | -23% | 0.0147 | 34 | Fig. 1E |
| **Experiment 4** |  |  |  |  |  |
| 1m *fog-2* | 10.5 ± 0.5 | -- | -- | 35 | Fig. S1A |
| 1f1m_3d *fog-2* | 6.6 ± 0.2 | -37% | <0.0001 | 33 | Fig. S1A |
| 2f1m_3d *fog-2* | 6.3 ± 0.2 | -40% | <0.0001 | 32 | Fig. S1A |
| 3f1m_3d *fog-2* | 6.4 ± 0.3 | -39% | <0.0001 | 31 | Fig. S1A |
| **Experiment 5** |  |  |  |  |  |
| 1m *fog-2* | 12.0 ± 0.4 | -- | -- | 40 | Fig. S1B |
| 1f1m_1.7d *fog-2* | 10.6 ± 0.5 | -12% | 0.0824 | 32 | Fig. S1B |
| 2f1m_1.7d *fog-2* | 9.7 ± 0.6 | -19% | 0.0435 | 37 | Fig. S1B |
| 3f1m_1.7d *fog-2* | 10.4 ± 0.6 | -13% | 0.1575 | 36 | Fig. S1B |
| 4f1m_1.7d *fog-2* | 9.3 ± 0.6 | -23% | 0.0041 | 36 | Fig. S1B |
| **Experiment 6** |  |  |  |  |  |
| 1m *fog-2* | 13.8 ± 0.7 | -- | -- | 35 | Fig. 1F |
| 3f1m_3d(D1-3) *fog-2* | 10.6 ± 0.8 | -23% | 0.0147 | 34 | Fig. 1F |
| 3f1m_3d(D6-8) *fog-2* | 10.8 ± 0.6 | -22% | 0.0022 | 37 | Fig. 1F |
| **Experiment 7** |  |  |  |  |  |
| 1m *fog-2* | 12.1 ± 0.6 | -- | -- | 32 | Fig. 2A |
| 1f1m_6d *daf-22* x *fog-2* | 9.0 ± 0.4 | -26% | 0.0001 | 29 | Fig. 2A |
| **Experiment 8** |  |  |  |  |  |
| 1m *daf-22* | 13.8 ± 0.6 | -- | -- | 40 | Fig. 2B |
| 1f1m_6d *fog-2* x *daf-22* | 7.4 ± 0.4 | -46% | <0.0001 | 34 | Fig. 2B |
| **Experiment 9** |  |  |  |  |  |
| 1m *fog-2* | 10.5 ± 0.5 | -- | -- | 35 | Fig. 2C |
| 3f1m_3d *fog-2* | 6.4 ± 0.3 | -39% | <0.0001 | 31 | Fig. 2C |
| 3f1m_3d+FUDR *fog-2* | 10.2 ± 0.4 | -3% | 0.7086 | 36 | Fig. 2C |
| **Experiment 10** |  |  |  |  |  |
| 1m *fog-2* | 13.8 ± 0.7 | -- | -- | 35 | Fig. S1C |
| 1f1m_6d *fog-2* | 10.3 ± 0.6 | -25% | 0.0006 | 31 | Fig. S1C |
| 1m+FUDR *fog-2* | 13.9 ± 0.4 | +1% | 0.4079 | 35 | Fig. S1C |
| 1f1m_6d+FUDR *fog-2* | 13.6 ± 0.5 | -1% | 0.3992 | 34 | Fig. S1C |
| 3f1m_3d *fog-2* | 10.6 ± 0.8 | -23% | 0.0147 | 34 | Fig. S1D |
| 3f1m_3d+FUDR *fog-2* | 14.3 ± 0.7 | +4% | 0.8740 | 32 | Fig. S1D |
| **Experiment 11 @26°C** |  |  |  |  |  |
| 1m *glp-1* | 8.0 ± 0.4 | -- | -- | 40 | Fig. 2D |
| 1f1m_6d *fog-2* x *glp-1* | 7.2 ± 0.4 | -10% | 0.3178 | 40 | Fig. 2D |
| **Experiment 12 @25°C** |  |  |  |  |  |
| 1m *glp-1* | 11.1 ± 1.0 | -- | -- | 27 | Fig. S1E |
| 1f1m_6d *fog-2* x *glp-1* | 11.1 ± 0.5 | 0% | 0.9149 | 43 | Fig. S1E |
| **Experiment 13 @26°C** |  |  |  |  |  |
| 1m *glp-1* | 9.6 ± 0.4 | -- | -- | 40 |  |
| 1f1m_6d *fog-2* x *glp-1* | 8.8 ± 0.5 | -8% | 0.238 | 40 |  |
| **Experiment 14** |  |  |  |  |  |
| 1m *pqm-1* | 11.9 ± 0.5 | -- | -- | 25 | Fig. 3E |
| 1f1m_6d *fog-2* x *pqm-1* | 11.0 ± 0.6 | -8% | 0.2782 | 29 | Fig. 3E |
| **Experiment 15** |  |  |  |  |  |
| 1m *fog-2* (L4440) | 12.6 ± 0.7 | -- | -- | 25 | Fig. 3F |
| 1f1m_4d *fog-2* (L4440) | 8.8 ± 0.5 | -30% | 0.0001 | 33 | Fig. 3F |
| 1m *fog-2* (*unc-62i*) | 11.9 ± 0.8 | -- | -- | 25 | Fig. 3F |
| 1f1m_4d *fog-2* (*unc-62i*) | 10.6 ± 0.5 | -11% | 0.1249 | 34 | Fig. 3F |
| **Experiment 16** |  |  |  |  |  |
| 1m *C. r.* | 31.4 ± 1.7 | -- | -- | 72 | Fig. 4A |
| 1f1m_6d *C. r.* | 15.7 ± 1.2 | -50% | <0.0001 | 28 | Fig. 4A |
| **Experiment 17** |  |  |  |  |  |
| 1m *C. e.* | 10.2 ± 0.6 | -- | -- | 35 | Fig. 4B |
| 1m1f_6d *C. e.* x *C. e.* | 7.4 ± 0.4 | -27% | 0.0001 | 35 | Fig. 4B |
| 1m1f_6d *C. r.* x *C. e.* | 7.4 ± 0.4 | -27% | 0.0003 | 35 | Fig. 4B |
| **Chinese Emperors LS** |  |  |  |  |  |
| Normal | 52.3 ± 1.0 | -- | -- | 234 | Fig. 7 |
| Promiscuous | 34.0 ± 1.9 | -35% | <0.0001 | 21 | Fig. 7 |
| **Experiment 18** |  |  |  |  |  |
| 1m *fog-2* | 12.0 ± 0.4 | -- | -- | 40 | Fig. 5A |
| 2m *fog-2* | 10.6 ± 0.4 | -12% | 0.0397 | 40 | Fig. 5A |
| 4m *fog-2* | 9.9 ± 0.4 | -18% | 0.0012 | 60 | Fig. 5A |
| 8m *fog-2* | 7.7 ± 0.2 | -36% | <0.0001 | 80 | Fig. 5A |
| **Experiment 19** |  |  |  |  |  |
| 1m+FUDR *fog-2* | 13.9 ± 0.4 | -- | -- | 35 | Fig. 5B |
| 8m+FUDR *fog-2* | 12.4 ± 0.3 | -11% | 0.0032 | 48 | Fig. 5B |
| 1m *fog-2* | 13.8 ± 0.7 | -- | -- | 35 | Fig. 5E |
| 8m *fog-2* | 9.8 ± 0.5 | -29% | <0.0001 | 48 | Fig. 5E |
| 8m *daf-22* | 14.7 ± 0.7 | +7% | 0.4039 | 48 | Fig. 5E |
| 1m+FUDR *daf-22* | 15.3 ± 0.3 | -- | -- | 35 | Fig. 5F |
| 8m+FUDR *daf-22* | 14.7 ± 0.3 | -4% | 0.2117 | 48 | Fig. 5F |
| 1m *daf-22* | 17.2 ± 0.6 | -- | -- | 35 |  |
| 8m *daf-22* | 14.7 ± 0.7 | -15% | 0.0660 | 48 |  |
| **Experiment 20** |  |  |  |  |  |
| 1m *daf-22* | 21.7 ± 1.2 | -- | -- | 32 | Fig. S4A |
| 8m *daf-22* | 18.8 ± 1.0 | -13% | 0.0394 | 38 | Fig. S4A |
| **Experiment 21** |  |  |  |  |  |
| 1m *C.r.* | 37.9 ± 1.1 | -- | -- | 120 | Fig. 5C |
| 8m *C.r.* | 31.0 ± 0.9 | -18% | <0.0001 | 160 | Fig. 5C |
| **Experiment 22** |  |  |  |  |  |
| 1m *C.r.* + FUDR | 30.8 ± 0.9 | -- | -- | 45 | Fig. 5D |
| 8m *C.r.* + FUDR | 30.8 ± 0.5 | 0% | 0.9217 | 112 | Fig. 5D |
| **Experiment 23** |  |  |  |  |  |
| 1m *daf-22* ctrl | 23.0 ± 0.9 | -- | -- | 30 | Fig. 5G |
| 1m *daf-22* ≤ 1mMCP | 17.3 ± 0.7 | -25% | <0.0001 | 29 | Fig. 5G |
| 1m *daf-22* ≤ 8mMCP | 16.1 ± 0.6 | -30% | <0.0001 | 30 | Fig. 5G |
| **Experiment 24** |  |  |  |  |  |
| *C.e.* herm ctrl | 14.4 ± 0.8 | -- | -- | 90 | Fig. 6A |
| *C.e.* herm ≤ *C.e.* MCP | 10.9 ± 0.6 | -24% | 0.0004 | 60 | Fig. 6A |
| *C.e.* herm ≤ *C.r.* MCP | 11.9 ± 0.5 | -17% | 0.0042 | 90 | Fig. 6A |
| *C.r.* fem ctrl | 15.8 ± 0.9 | -- | -- | 60 | Fig. 6B |
| *C.r.* fem ≤ *C.r.* MCP | 19.5 ± 1.3 | +23% | 0.0636 | 30 | Fig. 6B |
| *C.r.* fem ≤ *C.e.* MCP | 18.5 ± 0.9 | +17% | 0.1770 | 60 | Fig. 6B |
| **Experiment 25** |  |  |  |  |  |
| 1m *C. e.* | 12.1 ± 0.6 | -- | -- | 30 | Fig. 6C |
| 1m ≤ 8mMCP *C. e.* | 9.8 ± 0.4 | -19% | 0.0046 | 28 | Fig. 6C |
| 1f *C. e.* | 13.8 ± 0.7 | -- | -- | 30 | Fig. 6C |
| 1f ≤ 8mMCP *C. e.* | 12.6 ± 0.9 | -9% | 0.5965 | 29 | Fig. 6C |
| **Experiment 26** |  |  |  |  |  |
| 1m *C. r.* | 35.8 ± 2.0 | -- | -- | 34 | Fig. 6D |
| 1m ≤ 8mMCP *C. r.* | 37.8 ± 1.2 | +6% | 0.8501 | 34 | Fig. 6D |
| 1f *C. r.* | 27.6 ± 2.2 | -- | -- | 24 | Fig. 6D |
| 1f ≤ 8mMCP *C. r.* | 27.0 ± 2.5 | -2% | 0.8306 | 30 | Fig. 6D |

**Table S2.** Body size measurements.

| Genotype/<br>condition | N |  | Day 1 | Day 2 | Day 3 | Day 4 | Day 5 | Day 6 | Day 7 |
| --- | --- | --- | --- | --- | --- | --- | --- | --- | --- |
| *fog-2(q71)*<br>unmated | 31 | Body length ± SE (μm) | 903.6 ± 5.0 | 961.9 ± 5.6 | 982.8 ± 7.2 | 976.2 ± 7.2 | 959.0 ± 7.4 | 935.9 ± 8.1 | 931.5 ± 10.3 |
| *fog-2(q71)*<br>mated | 30 | | 893.8 ± 5.1 | 938.2 ± 5.6 | 955.4 ± 7.1 | 941.0 ± 9.5 | 913.7 ± 14.0 | 855.9 ± 19.4 | 851.4 ± 24.5 |
| @20°C |  | p value | 0.1813 | 0.004 | 0.0088 | 0.0044 | 0.0055 | 0.0004 | 0.0015 |
|  |  | % change | -1.1% | -2.5% | -2.8% | -3.6% | -4.7% | -8.6% | -8.6% |
| *glp-1(e2141)*<br>unmated | 40 | Body length ± SE (μm) | 789.6 ± 11.3 | 880.1 ± 6.4 | 894.0 ± 7.1 | 895.1 ± 7.3 | 888.8 ± 7.9 | 872.0 ± 8.8 | 874.3 ± 9.0 |
| *glp-1(e2141)*<br>mated | 40 | | 789.6 ± 11.3 | 887.0 ± 5.9 | 892.7 ± 8.1 | 894.0 ± 6.7 | 888.1 ± 10.8 | 868.6 ± 12.0 | 857.5 ± 11.4 |
| @26°C |  | p value | - | 0.4282 | 0.9083 | 0.9093 | 0.9567 | 0.8173 | 0.2510 |
|  |  | % change | - | 0.8% | -0.1% | -0.1% | -0.1% | -0.4% | -1.9% |

**Table S3.** Mated males microarray SAM rank table.

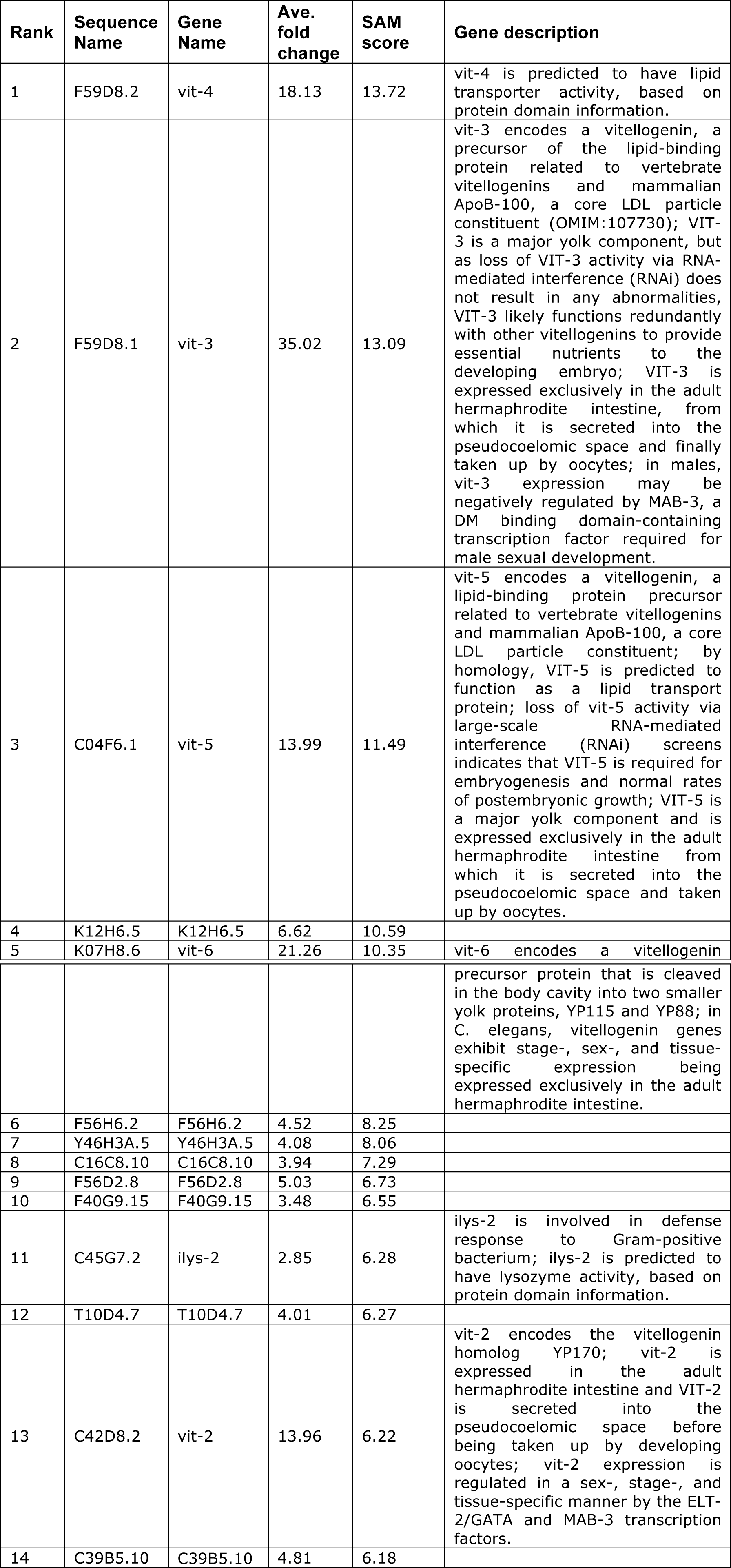
A. up-regulated genes (compared to unmated control)

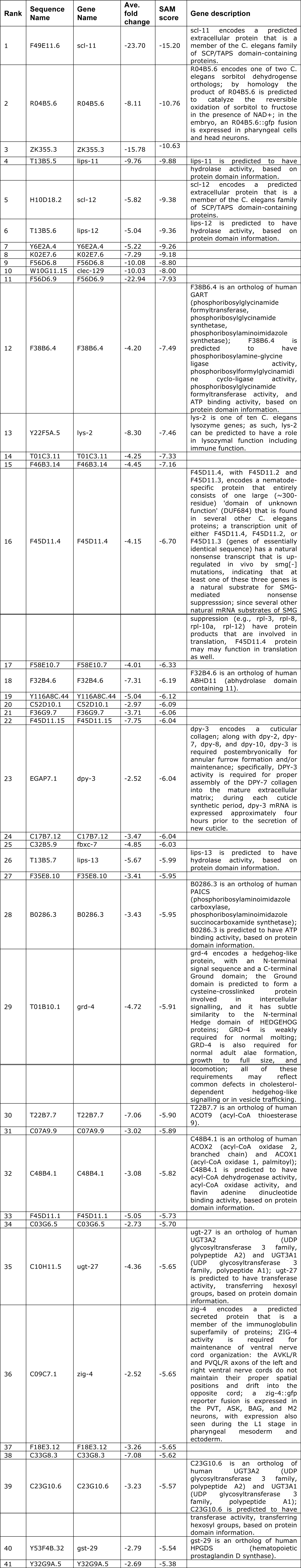
B. down-regulated genes (compared to unmated control)

**Table S4.**

List of Chinese Emperors.

